# Metabolic rewiring induces a viral mimicry-mediated quiescence in glioblastoma

**DOI:** 10.64898/2026.10.01.755088

**Authors:** Charles Chesnelong, Kinjal Desai, Connor S. Charton, Michelle Kushida, Chunying Yu, Paul Guilhamon, Lilian Lee, Sameer Agnihotri, Peter B. Dirks

## Abstract

Glioblastoma is a highly aggressive brain tumour driven by therapy-resistant glioblastoma stem cells (GSCs) that switch between dormant and proliferative states through poorly defined programs. In this study, we identify *SAT1*, encoding the rate-limiting polyamine-catabolic enzyme SSAT1, as enriched in quiescent GSCs, linking metabolic rewiring to dormancy. Pharmacologic induction of SSAT1 with the polyamine analogue N^1^,N^11^-Diethylnorspermine (DENSpm) triggers a chemoprotective, reversible growth arrest in patient-derived GSCs, independent of stemness, differentiation or senescence. SAT1- induced dormancy coincides with reduced eIF5A hypusination and global protein translation, and a marked rise in double-stranded RNA that activates type I interferon signalling. Knocking down the mitochondrial antiviral-signalling protein, essential for antiviral innate immunity, partially reverses this growth arrest, establishing viral mimicry as functionally required for dormancy. Single-cell and cohort analyses across GSCs and neural stem cells demonstrate that susceptibility to metabolically induced quiescence is set by an intrinsic inflammatory tone and lysosomal dependency, defining a context-dependent metabolic-immune axis which can be targeted to prevent GBM progression.

## Introduction

Glioblastoma (GBM) is the most aggressive and common type of primary malignant brain tumour in adults. Despite the initial effectiveness of surgery, radio- and chemotherapy, GBM invariably recurs, with <10% of patients surviving beyond 5 years.^1^ Brain tumour growth is sustained by a population of cancer stem cells at the apex of a functional cellular hierarchy, which can evade conventional therapy, repopulate the tumour, and seed recurrence.^2–6^ These cells exhibit long-term self-renewal, multilineage differentiation and high tumorigenic potential, features they share with the normal stem and progenitor cells of the tissue they arise from, such that brain tumour progression represents a distorted version of normal brain development or regeneration.^7–15^ Brain tumour stem cells can exist in a quiescent state or become activated and enter the cell cycle, with activation appearing to be a necessary step for downstream progenitor formation and tumour development.^16^ Selectively targeting glioblastoma stem cells (GSCs) is therefore essential to improve clinical outcomes, yet the GSC properties that drive tumour growth, and the potentially targetable regulators governing the switch between quiescent and activated states remain poorly understood.

Within the phenotypically distinct tumour compartments, the largely quiescent GSC population is itself heterogeneous; reflecting intrinsic biological diversity that may arise from distinct cells of origin, environmental context or precursor state. Analyses of their molecular and functional characteristics have resolved this heterogeneity into defined subgroups.^17–23^ From these studies, two distinct GSC populations have emerged: one that displays classical stemness properties and is associated with a proneural phenotype, and a second, more aggressive population that displays features more associated with a mesenchymal phenotype. Single-cell transcriptomics has since refined this perspective, and in our view, placed GSCs along a continuous transcriptional gradient between two states reminiscent of normal neural development, or an inflammatory injury response.^24^ GSCs distributed across this spectrum possess distinct characteristics and likely unique exploitable vulnerabilities, although whether these states are fully interconvertible is not clear.

In this study, we examine quiescence as a core stem cell feature central to the role of GSCs in tumour growth and therapeutic resistance, and identify *SAT1* as a regulator of GSC quiescence. *SAT1* encodes the rate-limiting enzyme of polyamine catabolism, SSAT1, and its overexpression markedly depletes two key polyamine pools, spermidine (Spd) and spermine (Spm). Polyamines, which are naturally occurring polycations essential for the proliferation of eukaryotic cells, decline with age, and their supplementation extends lifespan.^25–29^ Notably, they are elevated in cancer cells, thus presenting as attractive therapeutic targets,^30^ and have been implicated in inflammatory responses.^31^ Polyamine regulation is also central to the response to pathogens,^31,32^ with *SAT1* itself being a bona fide interferon-stimulated gene (ISG) with a defined role in antiviral defence.^33^ Our results indicate that *SAT1* promotes a viral mimicry-driven quiescent state, with developmental GSCs being particularly sensitive to induced quiescence.

Interestingly, polyamine depletion-induced quiescence was accompanied by upregulation of TFEB, a regulator of innate immunity and lysosome biogenesis, implicating altered lysosomal function as a feature of this stem cell state. Combining pharmacological inhibition of autophagosome–lysosome fusion with DENSpm in vitro prevented GSCs from resuming growth after polyamine depletion, indicating that lysosomal function sustains the reversibility of induced quiescence and identifying it as a targetable dependency in GBM.

## Results

### Identification of quiescent GSCs in culture

We first sought to determine if quiescent sub-populations existed in patient-derived glioblastoma stem cells (GSCs). We probed single-cell RNA sequencing (scRNAseq) performed on a large cohort of GSC cultures derived from patient tumours. By stratifying the samples for cycling vs. non-cycling cells based on G1/S and G2/M cell cycle scoring, we identified quiescent populations in every GSC sample, with frequencies ranging from <1 to 45% (Fig. 1A–D). Utilising previously published gene signatures of quiescence in neural stem cells (Neural.G0)^34^ and glioblastoma (GBM) (Q118),^6^ we validated that the non-cycling population we had identified is enriched in quiescent cells (Supplementary Fig. 1A–D).

**Figure 1:**
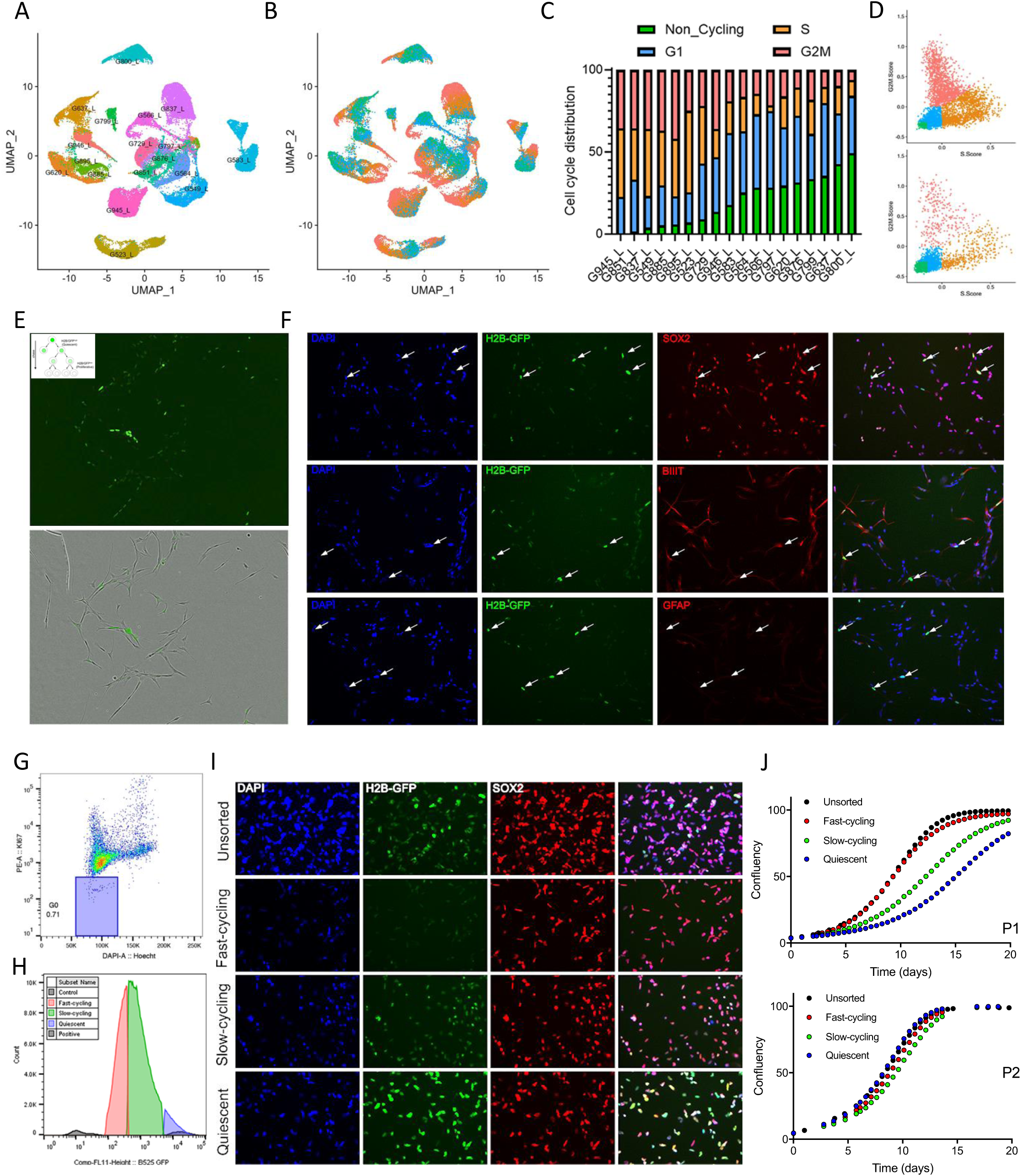
Identification of quiescent GSCs in culture **(A)** scRNAseq UMAP colored based on GSCs IDs. **(B)** UMAP recolored according to cell cycle phases. **(C–D)** Bar graph representing cell distribution in the distinct cell cycle phases (C) and representative cell cycle scatter plots (D). **(E)** Representative images of label retaining cells. **(F)** Immunofluorescence images of GSCs stained for SOX2, Beta III Tubulin, and GFAP. **(G)** Representative flow cytometry figures illustrating the presence of a quiescent G0 population in GSCs (G564). **(H)** Flow cytometry graphs illustrating the sorting of quiescent, slow-cycling, and fast-cycling cells based on H2B-GFP intensity. **(I)** Representative images with SOX2 immunostaining. **(J)** Graph showing the viability of the three sorted populations (P1 and P2) after plating post-sorting across 20 days. Data presented in E-J were acquired with G549.

To functionally corroborate the presence of the quiescent sub-populations within the GSC cultures, we performed a label retention experiment using a fluorescently tagged H2B histone protein. This method allows for stable incorporation of the fluorescent tag into nucleosomes, providing a persistent signal that is only diluted when a labelled cell undergoes division. Despite the presence of mitogens in cell culture media, we detected a Ki67-negative label retaining population (Fig. 1E–F). These cells express the stem cell marker *SOX2* but not markers of astrocytic or neuronal differentiation, GFAP and βIII-Tubulin (Fig. 1F). Importantly, when isolated, these slow-cycling cells readily resume growth and recapitulate the growth rate of the parental culture upon serial passaging (Fig. 1G-J; Supplementary Fig. 1E–F).

### *SAT1* is overexpressed in non-cycling GSCs

To study the pathways underlying cellular dormancy, we probed the scRNAseq dataset for genes upregulated in the quiescent cell population (Fig. 2A–B). Gene Ontology terms enriched in these cells across cell lines were largely related to immune and inflammatory processes (Fig. 2B, Supplementary Fig. 2A, Supplementary Tables 2 and 3); signatures typically associated with the injury response GSC transcriptional profile.

**Figure 2:**
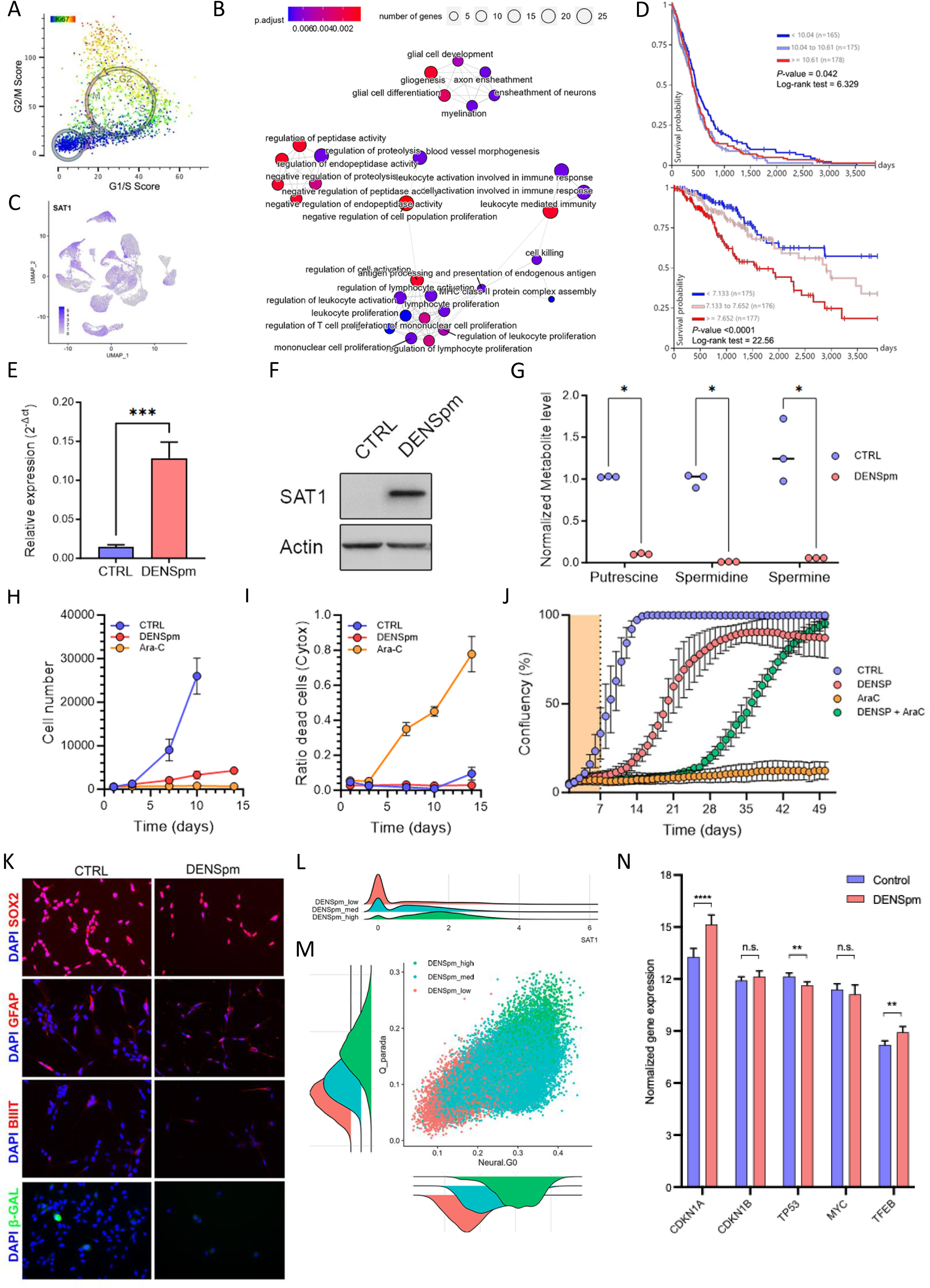
SAT1 regulates a quiescent phenotype in GSCs **(A)** scRNAseq cells stratified on G1/S and G2M scores, coloured by Ki67 expression. **(B)** Emapplot of significantly enriched GO terms in non-cycling GSCs from scRNAseq data. **(C)** SAT1 feature plot. **(D)** Kaplan-Meier survival curves according to SAT1 expression in GBM (top) and low grade glioma (bottom). **(E)** Bar graph representing SAT1 expression as measured by RT-qPCR and **(F)** Western blot. **(G)** Plot showing polyamine levels (G549, G564, G583) measured by mass spectrometry and graphs showing **(H)** GSCs growth and **(I)** cell death upon DENSpm (10μM) or Ara-C (1μM) treatment (G549). **(J)** Graph showing GSCs growth with and without DENSpm and/or Ara-C treatment followed by a wash after a 7-day treatment (G549). **(K)** Representative immunofluorescence images of GSCs treated or not with DENSpm (7 days, 10μM) and stained for SOX2 (stem marker), GFAP (astrocytic marker), and Beta III Tubulin (neuronal marker) to evaluate differentiation, as well as Beta-Galactosidase activity to assess senescence. **(L)** SAT1 ridge plot of cells segregated according to DENSpm_Q and **(M)** Quiescence scatter plots (NSC: Neural.G0 and GBM: Q118) colored based on DENSpm_Q. All data presented in E– K was acquired using G549. **(N)** Bar graph representing normalized gene expression as measured by bulk RNA sequencing between vehicle (control) and DENSpm treated GSCs.

Differential expression analysis further revealed that SAT1 was consistently overexpressed in the quiescent GSC population (Fig. 2C, Supplementary Fig. 2B, Supplementary Table 2). *SAT1* encodes spermidine/spermine N1-acetyltransferase 1 (SSAT1), the rate-limiting enzyme of polyamine catabolism, which acetylates spermine (Spm) and spermidine (Spd) to trigger their export and depletion. Polyamines are polycationic metabolites essential for proliferation: they are required for cell-cycle progression, stabilise nucleic acids and chromatin, and in the case of spermidine, serve as the sole donor for hypusination of the translation factor eIF5A. Their depletion therefore may impair growth through several mechanisms, and *SAT1* upregulation is accordingly known to suppress translation and induce growth arrest in both normal and cancer contexts. While high *SAT1* expression promotes growth arrest, it is associated with shorter overall survival in low-grade gliomas and GBMs (Fig. 2D). This apparent paradox points to a role for SAT1 not in eliminating tumour cells but in potentially maintaining a quiescent stem cell pool that survives treatment and drives recurrence. We therefore hypothesized that *SAT1* enforces quiescence by depleting spermine and spermidine, sustaining a dormant, therapy-resistant reservoir.

### *SAT1* regulates a quiescent phenotype in GSCs

To investigate induction of the quiescent phenotype in GSCs, we recapitulated *SAT1* upregulation using the polyamine analogue N^1^,N^11^-Diethylnorspermine (DENSpm). DENSpm treatment induced *SAT1* expression and SSAT1 protein abundance (Fig. 2E–F) and triggered polyamine depletion (Fig. 2G). It was strongly cytostatic, with minimal associated cell death (Fig. 2H–I, Supplementary Fig. 2C), and produced a long-term but reversible growth arrest (Fig. 2J) without altering the expression of stem, differentiation, or senescence-associated markers (Fig. 2K). Critically, cells concurrently treated with the anti-mitotic agent Cytarabine (AraC) and DENSpm markedly reduced cell death relative to vehicle-treated controls (Fig. 2J), establishing that SAT1-induced dormancy confers chemoprotection. Limiting dilution assays showed that although proliferation was impaired during treatment, self-renewal capacity was retained following treatment (Supplementary Fig. 2D–E), indicating that the arrest suppresses growth without exhausting the stem cell pool.

We next performed bulk RNA sequencing and derived a DENSpm-induced quiescence signature (DENSpm_Q score) from the genes significantly upregulated after one week of DENSpm treatment (Supplementary Fig, 2F, Supplementary Tables 1 and 4). Applied to our scRNAseq data, this score was enriched in non-cycling cells (Supplementary Fig. 2G–H). Importantly, cells with a high DENSpm_Q score also overexpressed *SAT1* (Fig. 2L) and showed elevated Neural.G0 and Q118 scores (Fig. 2M), validating the signature. Among the genes upregulated in DENSpm-treated GSCs was TFEB, a regulator of innate immunity and lysosome biogenesis, as well as *CDKN1A*, which encodes the cyclin-dependent kinase inhibitor p21 (Fig. 2N), and whose increased expression likely contributes to the transcriptional basis of the growth arrest.

### DENSpm triggers translation arrest and the activation of antiviral response pathways

The dynamic maintenance of a balanced, functional proteome, or proteostasis, is a defining property of dormant cells, allowing them to persist without responding to every environmental cue. Depletion of polyamines is known to impair translation, and indeed, spermidine is the only known substrate for the hypusination and activation of eukaryotic translation initiation factor 5A (eIF5A). Consistent with this, both *SAT1* overexpression and DENSpm treatment sharply reduced eIF5A hypusination, and its activation (Supplementary Fig. 3A). Using a puromycin analogue to detect translational activity, we also determined that DENSpm impairs translation in GSCs (Fig. 3A–B).

**Figure 3:**
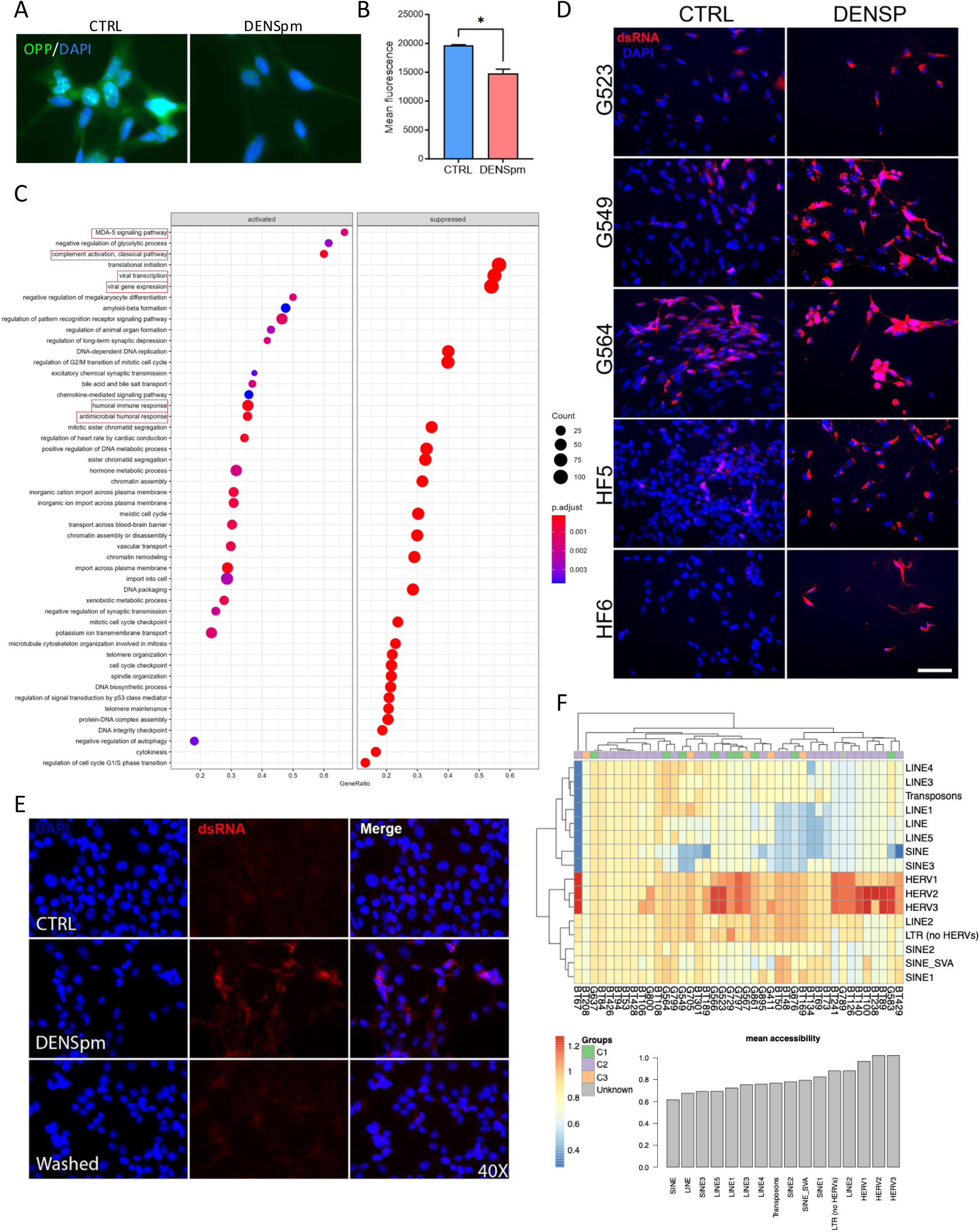
DENSpm triggers translation arrest and the activation of antiviral response pathways **(A)** Representative images of O-propargyl-puromycin (OPP) labeling to assess total protein synthesis and **(B)** bar graph of quantified data. **(C)** Graph representing activated and suppressed pathways identified through GSEA in GSCs upon DENSpm treatment. Pathways related to pathogen response are outlined in red. **(D)** Representative images of dsRNA immunolabelling (J2 antibody) in GSCs and Human fetal NSCs treated for a week with DENSpm, and **(E)** post DENSpm wash (G549). **(F)** Heatmap and bar graph showing mean accessibility across diverse families of transposable elements. Data derived from ATAC-seq performed on a large collection of untreated GSCs.^39^

Next, we evaluated the role of *SAT1* as an epigenetic modifier acting through S-Adenosyl Methionine (SAM). While SAM is well known for its role as a methyl donor to various methyltransferases, it is also essential for polyamine biosynthesis and was shown to be depleted upon DENSpm treatment due to an increased flux through the polyamine metabolic pathway.^35^ We also found that low polyamines level correlates with low S-Adenosyl-methionine (SAM) and DNA hypomethylation in untreated GSCs and hfNSCs (Supplementary Fig. 3B). We therefore reasoned that SAM depletion might drive epigenetic changes underlying the quiescent phenotype. Although DENSpm treatment depleted SAM (Supplementary Fig. 3C), it did not globally alter DNA methylation (Supplementary Fig. 3D–F) or cause appreciable global histone demethylation (Supplementary Fig. 3G), suggesting that focal, alternative pathways are responsible for the induction of quiescence triggered by DENSpm treatment.

To identify the molecular programs underlying the DENSpm-induced growth arrest, we performed gene set enrichment analysis on RNA sequencing from DENSpm-treated GSCs. As expected, cell cycle pathways were suppressed; conversely, pathways for the detection of cytoplasmic double-stranded RNA (dsRNA), along with other pathogen-defence, immune and inflammatory programs, were significantly activated upon treatment (Fig. 3C, Supplementary Table 5). This echoed our initial GO analysis of the scRNAseq data, which had highlighted immune activation in the non-cycling compartment (Fig. 2B).

Defence-response-to-virus, type I interferon pathways were likewise activated upon SAT1 overexpression (Supplementary Fig. 3H–K, Supplementary Tables 6 and 7). We therefore investigated whether *SAT1* drives this antiviral response. By immunofluorescence, we observed a marked increase in dsRNA immunoreactivity upon treatment across GSCs and neural stem cells (NSCs) (Fig. 3D); this increase was reversible when DENSpm was washed out after seven days (Fig. 3E, Supplementary Fig. 4A). We postulated that *SAT1* upregulation drives a viral mimicry process that maintains cells in a quiescent state. Viral mimicry refers to the endogenous expression of dsRNA, often from reactivated endogenous retroviral elements (ERVs), which triggers an antiviral response, leading to production of type I and III interferons (IFN I/III) that act in autocrine and paracrine fashion to induce interferon-stimulated genes (ISGs). ISGs restrict viral replication but also constrain cellular replication and translation. Notably, *SAT1* is itself a bona fide ISG, which could establish a positive feedback loop that reinforces quiescence in GSCs.^33^

ERV loci are more accessible in immature haematopoietic stem cells than in their differentiated progeny in the blood system.^36^ Consistently, in GSCs, we found ERVs to be the most accessible transposable element subfamily (Fig. 3F, Supplementary Fig. 4B–C), suggesting that these cells harbour epigenetically active ERVs whose abundance may be further tuned transcriptionally and post-transcriptionally, a process in which polyamines themselves may influence dsRNA stability, detection and degradation.^37^

### Viral mimicry and interferon signalling mediate DENSpm-induced quiescence

While the regulatory mechanisms controlling ERVs upregulation remains unclear, we showed in untreated GSCs, that ERVs expression correlates with the expression of IFNβ, the main antiviral response mediator and downstream ISGs (Supplementary Fig. 4D–E). This finding highlights a spectrum of activation of an interferon mediated antiviral response amongst GSCs. Remarkably, while *SAT1* RNA levels were not predictive, GSCs with high levels of viral mimicry, as characterized by high ERV and ISG z-scores, have the highest SSAT1 protein level (Supplementary Fig. 4D, F–G). Importantly, we showed activation of a viral mimicry response upon DENSpm treatment by confirming increased expression of ERVs, IFN type I/III and ISGs (Fig. 4A–C, Supplementary Fig. 4H). Activation of a dsRNA specific antiviral response was also supported by increased phosphorylation and nuclear translocation of IRF7 upon treatment (Fig. 4D–E).

**Figure 4:**
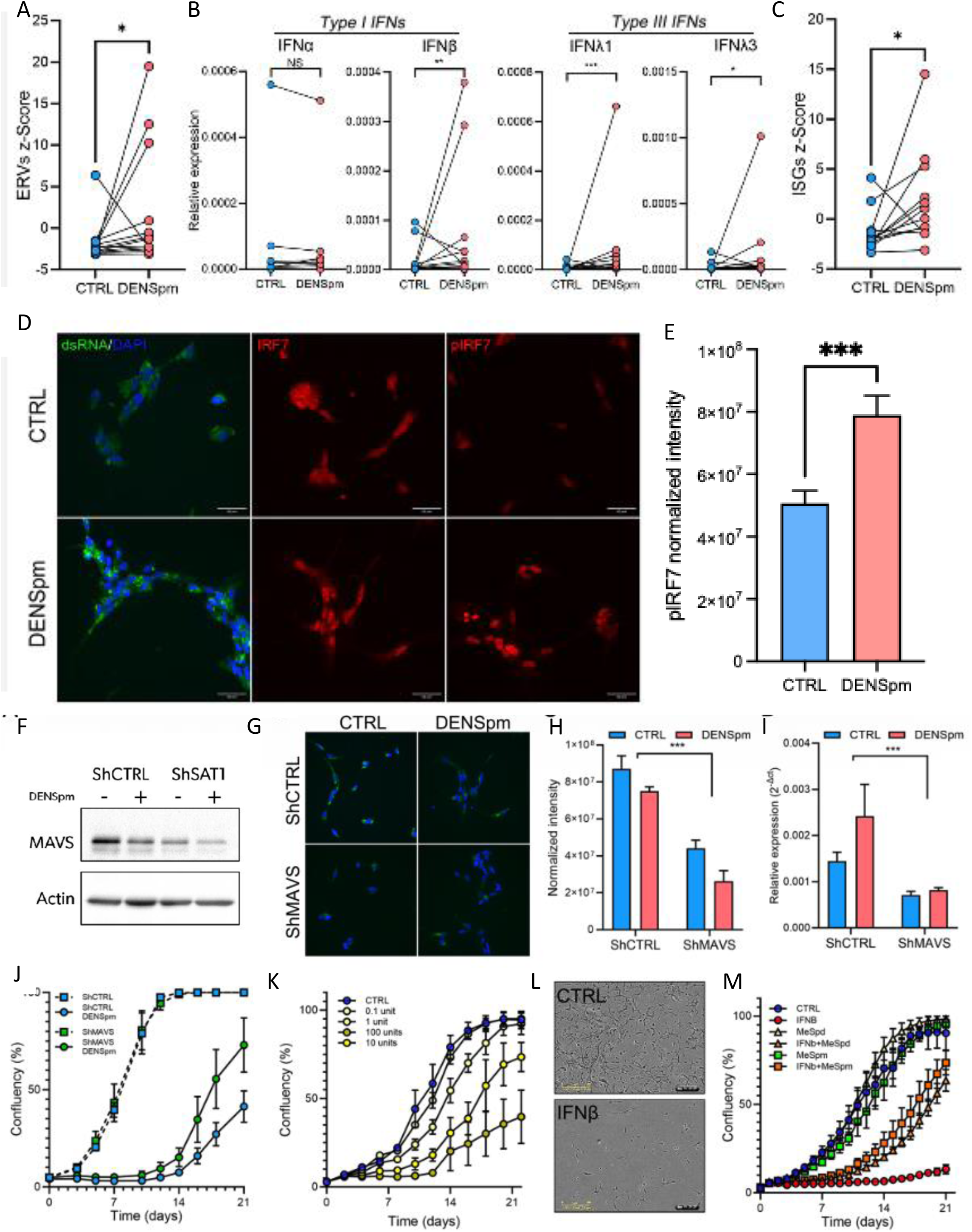
Viral mimicry and interferon signalling mediate DENSpm-induced quiescence **(A)** Aligned dot plots of ERVs z-score, **(B)** relative expression of type I (IFNα, IFNβ) and type III (IFNλ1 and IFNλ3) interferons, and **(C)** Aligned dot plots of ISGs z-score. **(D)** Representative immunofluorescence images of dsRNA labelling (J2 antibody), IRF7, phosphorylated IRF7 and **(E)** quantification of pIRF7 immunofluorescence. **(F)** Western blot, **(G)** representative immunofluorescence images, **(H)** bar graph of fluorescence signal quantification, and **(I)** relative expression of MAVS to assess knock-down efficiency. **(J)** Growth curves showing the impact of MAVS knock-down on DENSpm induced quiescence (7-day treatment course). **(K)** Growth curves of GSC treated with IFNβ. **(L)** Representative images of IFNβ impact on GSC growth as well as morphology (100 units IFNβ), and **(M)** rescue experiment with methylated polyamine analogues^38^ (1000 units IFNβ). All experiments performed with G523.

We next determined the role of Mitochondrial Antiviral Signaling Protein (MAVS), an essential mediator of the dsRNA dependent antiviral response, in GSCs. By MAVS knockdown, we partially rescued DENSpm-induced growth arrest validating a role for viral mimicry in this quiescent phenotype (Fig. 4F– J). IFNβ importantly induces a similar cytostatic effect to that of DENSpm in GSCs (Fig. 4K–L).

Methylated polyamine analogues, which cannot be acetylated by SAT1,^38^ were able to rescue the induction of this phenotype, together demonstrating how polyamine metabolism and viral mimicry are central to the induction of this quiescent state (Fig. 4M).

### DENSpm response is dependent on basal viral mimicry state

Systematic characterisation of DENSpm-induced quiescence in our cohort of GSCs and NSCs, revealed three distinct DENSpm response subgroups distinguished by whether cells entered the quiescent state and, having entered, whether they could leave it: non-reversible, reversible, and resistant (Fig. 5A and Supplementary Fig. 5A). Resistant GSCs continued to proliferate through treatment despite some impact on growth. Of the lines that did arrest, reversible GSCs readily re-entered the cell cycle once treatment was withdrawn, whereas non-reversible GSCs entered a deep quiescent state from which they did not recover. These outcomes tracked with the strength of the viral mimicry response. Induction of ERVs, IFN and ISGs upon DENSpm was stronger in non-reversible than in reversible GSCs and, more strikingly, these programs were globally downregulated upon treatment in most resistant lines (Fig. 5B, Supplementary Fig. 5B–C). (Fig. 5B, Supplementary Fig. 5B–C).

**Figure 5:**
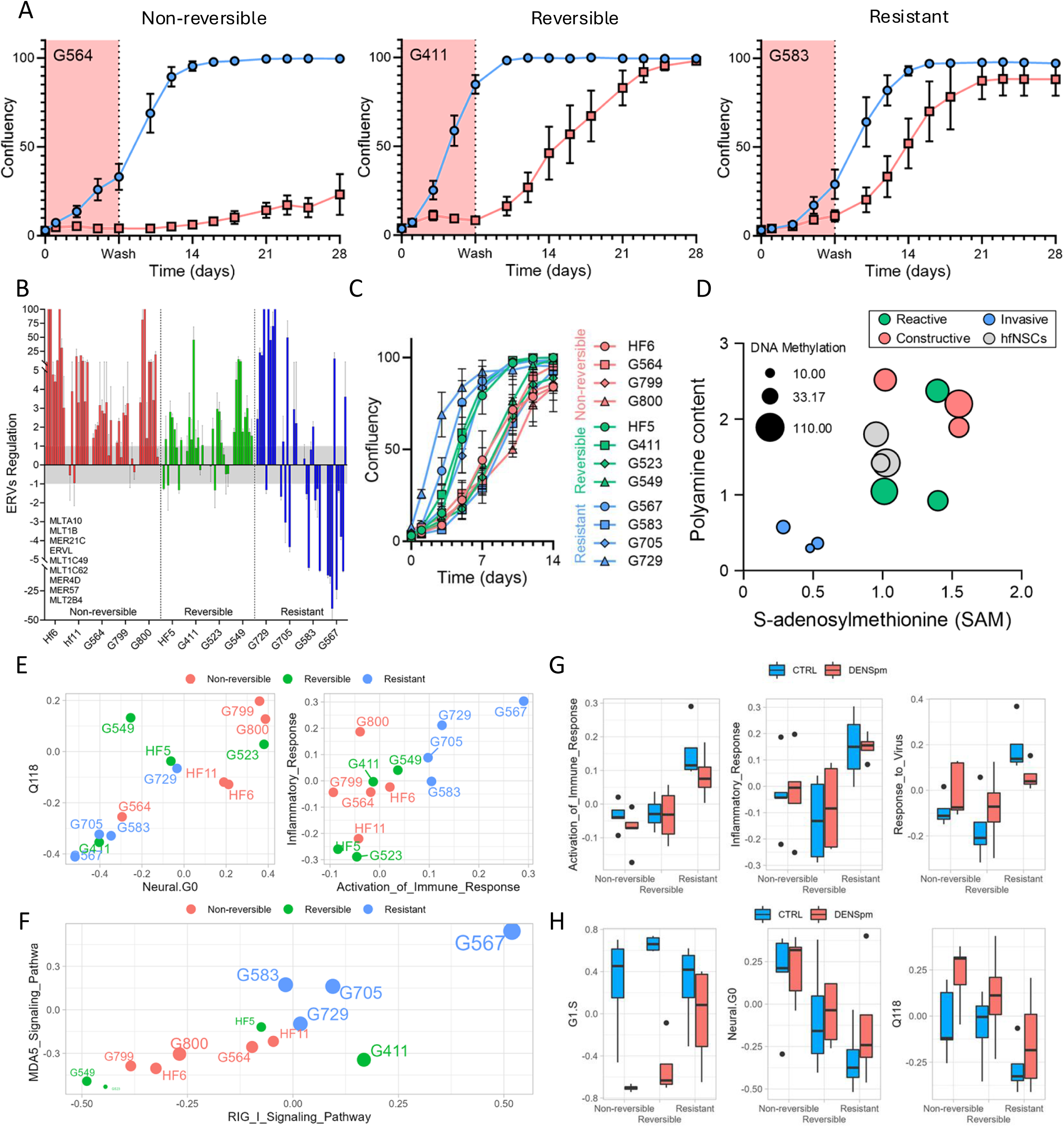
GSC inflammatory state dictates DENSpm response **(A)** Growth curves of GSCs and NSCs upon a 7-day DENSpm treatment (pink shaded area) followed by a wash (blue: CTRL; red: DENSpm). GSCs and NSCs are segregated according to DENSpm response; Non-reversible, Reversible, Resistant. **(B)** Bar graph of ERVs regulation after a 7-day DENSpm treatment and measured by RT-qPCR in GSCs and NSCs segregated according to response subgroups. **(C)** Basal growth curves of GSCs and NSCs colored according to DENSpm response groups. **(D)** Scatter plot of GSCs polyamine content, SAM level, and global DNA methylation (data used in Supplementary Fig. 2B) coloured according to recent GSC classification.^39^ **(E)** Scatter plots of untreated GSCs colored by DENSpm response and scattered according to the activity of quiescence signatures (Q118 and Neural.G0), activity inflammatory pathways, **(F)** dsRNA signaling pathways, and viral response pathway. **(G)** Box graphs of inflammatory and virus response pathways activity as well as **(H)** activity of cycling and quiescence pathways in the distinct GSCs response groups GSCs upon DENSpm treatment. Data presented E–H is based on bulk RNA sequencing of 13 GSC and NSC lines treated with DENSpm or vehicle.

The DENSpm response subgroups also match GSC growth characteristics. Non-reversible GSCs are typically slower-growing cells with a larger fraction of quiescent cells while resistant ones are more proliferative (Fig. 1C, 5C). Globally, DENSpm responses also align with the GSCs’ position along the developmental to injury response transcriptional axis.^24^ Resistant GSCs appear enriched in a particular subtype that we recently distinguished amongst more injury response GSCs; invasive GSCs (Supplementary Fig. 5D).^39^ This group has a similar transcriptomic profile as the reactive profile but presents a distinct epigenetic state, with more invasive behaviour and an association with lower survival in patients. While we have shown previously that these GSCs are globally hypomethylated, we now add that they are also characterized by lower polyamine and SAM content (Fig. 5D). Consequently, these GSCs may be acclimated to low polyamine level, possibly underlying their resistance to DENSpm.

Moreover, putative alterations of the polyamine metabolic pathway in invasive GSCs may result in SAM depletion, promote DNA hypomethylation, and subsequently enhance ERV reactivation. A sustained viral mimicry state, or a form of tolerance arising from chronically engaged innate immune signalling, could explain how these cells develop resistance to these inflammatory pathways, either by adaptation or selection.

### GSCs inflammatory state dictates DENSpm response

We found that DENSpm response tightly correlates with transcriptional profiles that reflect the quiescent and inflammatory state (Fig. 5E). This coupling of inflammatory and quiescent programs is reminiscent of normal human haematopoietic stem cells retaining memory of prior inflammatory stress.^40^ Resistance to DENSpm-induced quiescence is in fact predicted by high baseline activity of dsRNA sensing (measured by *MDA5* and *RIG-1* transcript levels) and defense response to virus pathways (Fig. 5F–G, Supplementary Fig. 5E–F, Supplementary Table 8). Basal activity of inflammatory and immune responses are elevated in resistant GSCs (Fig. 5E,G). Notably, these pathways remain mostly unaffected upon DENSpm treatment (Fig. 5G). However, the activity of defense response to virus pathway, is specifically upregulated upon DENSpm and most significantly so in non-reversible GSCs (Fig. 5G, Supplementary Fig. 5G–H, and Supplementary Table 9). Strikingly, resistant GSCs significantly down- regulate this defense response to virus pathway upon DENSpm (Fig. 5G, Supplementary Fig. 5G–H, and Supplementary Table 9). We further confirm that cell cycle programs are repressed upon DENSpm treatment in reversible and non-reversible GSCs while barely impacted in resistant GSCs (Fig. 5H). More importantly, we also show that DENSpm response groups differ on their basal quiescent state based on Neural.G0 and Q118 signatures and confirm upregulation of the GBM quiescent signature (Q118) upon DENSpm treatment (Fig. 5E,H). These results tightly link DENSpm response with basal overall inflammatory state, induction of antiviral response and quiescence in GSCs. Our results suggest that a viral mimicry-driven quiescence is specific to slower-growing, stem-like, neuro-developmental GSCs while more proliferative, inflammatory-poised progenitor-like GSCs are impervious to these additional cues.

We next asked whether non-reversible, reversible and resistant GSCs represent distinct entities coexisting within GSC cultures. To address this, we derived signatures for each group from the genes exclusively upregulated by DENSpm in that group (Supplementary Fig. 6A). Applied to our bulk transcriptomic database, the non-reversible and reversible scores marked more quiescent GSCs (Supplementary Fig. 6B– C), while the resistant scores are more associated with an injury response transcriptional state (Supplementary Fig. 6D–E). Because quiescent cells form only a small subset of any GSC culture, we turned to our single-cell data to resolve these populations directly. The three signatures distinguished cells that differed in cycling state, quiescence and developmental/injury-response identity (Supplementary Fig. 6F): non-reversible cells were the most quiescent (Q118, Neural.G0), whereas resistant cells were more proliferative and more injury response-like (Supplementary Fig. 6F).

### Lysosomal biogenesis is an exploitable vulnerability of quiescent cells

The three GSC response groups are distinguishable by polyamine content. Putrescine in particular is highest in non-reversible and lowest in resistant GSCs (Fig. 6A). This pattern suggests that non-reversible GSCs are especially reliant on polyamines, which may account for their extreme sensitivity to DENSpm. Consistent with this, polyamine content correlates with quiescence (Neural.G0), inflammatory response, and defense-response-to-virus signatures (Fig. 6B).

**Figure 6:**
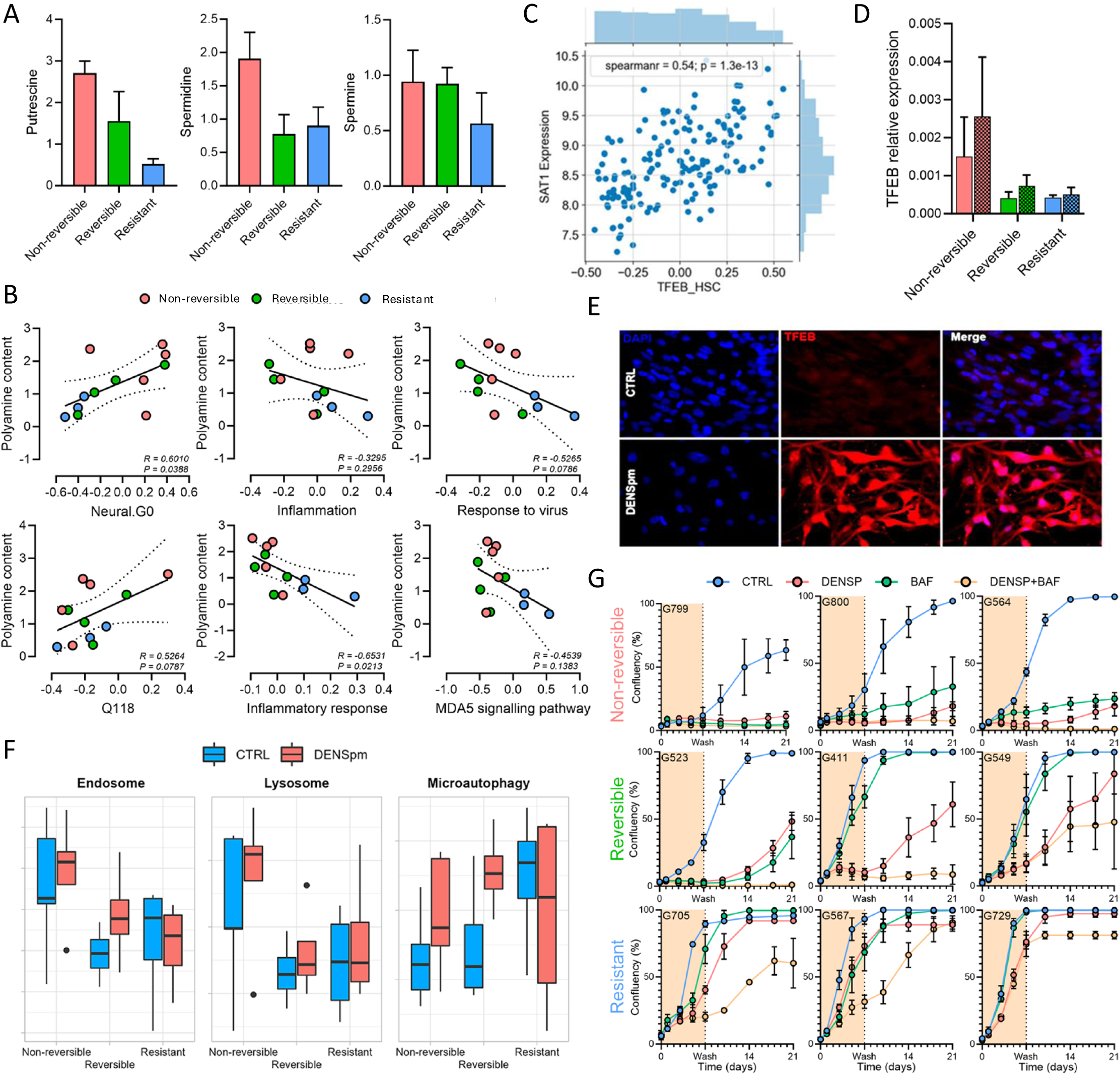
Quiescent cells are vulnerable to lysosome pathway disruption **(A)** Bar graphs of polyamine content in the GSCs DENSpm response groups. **(B)** Scatter plots with Pearson correlation of polyamine content with quiescence, inflammation, and virus response pathways in GSCs colored according to DENSpm response. **(C)** Scatter plot with Spearman correlation of TCGA GBM samples according to SAT1 expression and TFEB_HSC signature. **(D)** Bar graph of TFEB expression upon DENSpm measured by RT-qPCR in GSCs grouped according to DENSpm response and **(E)** representative TFEB immunofluorescence images (G564). **(F)** Box graphs of lysosome biogenesis related gene ontology pathways in GSCs treated with DENSpm and grouped according to DENSpm response. **(G)** Growth curves of GSCs upon a 7-day DENSpm and/or bafilomycin treatment. Graphs are arranged according to DENSpm response groups (vertically) and DENSpm sensitivity within groups (horizontally).

Cellular polyamine levels are tightly controlled through the balance of biosynthesis, catabolism, export and import, yet the mechanisms of polyamine uptake remain poorly defined. The lysosomal pathway was recently shown to contribute substantially to total cellular polyamine content. Polyamines are internalised by endocytosis and transported into the cytosol by the late endo-lysosomal transporter ATP13A2.^41,42^ ATP13A2, and potentially other P5B-ATPases,^43,44^ may therefore shape the DENSpm response (Supplementary Fig. 6G). Beyond uptake, these transporters prevent lysosomal polyamine accumulation and are required to maintain lysosomal integrity and function.^41,45^

Thus, polyamine handling is linked to lysosomal function, and in turn to quiescence. Lysosomes are central to autophagy, a process on which quiescent cells particularly depend, and TFEB, a master regulator of lysosome biogenesis and autophagy, is sufficient to induce quiescence in haematopoietic stem cells.^46^ Several lines of evidence place this axis in our system. The TFEB-induced quiescence signature correlates with SAT1 expression in glioma (Fig. 6C); TFEB levels are elevated in non-reversible GSCs and are strongly induced by DENSpm at both the RNA and protein level (Fig. 6D–E); and endosomal, lysosomal and lysosomal micro-autophagy genes are likewise elevated in non-reversible GSCs and upregulated upon DENSpm treatment (Fig. 6F).

We therefore reasoned that non-reversible GSCs are especially sensitive to polyamine depletion and consequently dependent on the lysosomal pathway to sustain their polyamine pools. To test this dependency, and a potential synergy with DENSpm, we used bafilomycin, an inhibitor of the vacuolar H+-ATPase and of autophagosome–lysosome fusion.^47^ Non-reversible GSCs were highly sensitive to bafilomycin, whereas resistant GSCs were largely refractory (Fig. 6G). In reversible GSCs, bafilomycin alone had only a minor effect but markedly impaired recovery following DENSpm treatment (Fig. 6G).

Taken together, these results define a metabolic–immune circuit governing GSC dormancy: SAT1-driven polyamine catabolism engages a viral mimicry response that enforces a reversible, chemoprotective quiescent state, the maintenance of which depends on lysosomal handling of polyamines. Susceptibility to this program is not uniform but stratifies GSCs into distinct response groups, set by their intrinsic inflammatory tone and lysosomal dependency. In the GSCs most reliant on this circuit, its disruption by blocking lysosomal function during or after DENSpm treatment prevents the recovery that would otherwise regenerate the tumour. These findings expose the dependencies of the dormant state as a targetable vulnerability in the cell population that drives GBM recurrence.

## Discussion

We demonstrate here that SAT1 promotes a viral mimicry-driven quiescent state in slower-growing, stem-like GSCs that depend on the lysosomal pathway for polyamine uptake. More proliferative, inflammation-poised progenitor-like GSCs, by contrast, appear already adapted to perturbation of these pathways. These cells also carry low basal polyamine levels, raising the possibility that long-term adaptation to polyamine depletion drives a pro-inflammatory state that is refractory to quiescence and may underlie the mesenchymal shift frequently observed in GBM at recurrence.^48^ Together, our findings support a model in which polyamine metabolism, viral mimicry, lysosomal biogenesis, and the inflammatory context of the tumour converge to regulate GSC quiescence.

The three response groups represent varying levels of innate competence set by basal innate immune tone to enter the dormant state. GSCs with a chronically engaged viral mimicry pathway are more resistant to DENSpm-induced quiescence, whereas GSCs with low basal tone retain the capacity to respond.

Quiescence itself remains a notoriously difficult cancer stem cell property to target directly. Our work defines such cell states in GBM and identifies a metabolic–immune circuit, SAT1-driven polyamine catabolism feeding a MAVS-dependent viral mimicry response, coupled to a lysosomal dependency, that maintains the quiescent stem cell state. These dependencies represent actionable vulnerabilities. A comparable phenomenon has recently been described in normal human haematopoiesis, where a subset of HSCs retains a molecular memory of prior inflammatory stress and adopts a quiescent state with restrained output,^40^ suggesting that the circuit we describe is not a tumour-specific adaptation but a conserved stem cell program that GBM retains and exploits.

Three approaches to engaging stem cell quiescence emerge. One is to promote quiescence exit and proliferation to sensitise cells to conventional agents, as achieved in chronic myelogenous leukaemia by inhibiting pro-survival regulators to render otherwise dormant stem cells susceptible to tyrosine kinase inhibition or cytotoxic ablation.^49^ Another, as established in SHH-subgroup medulloblastoma, is to constrain the quiescent stem cell state and block its proliferative activation as a necessary step in tumorigenesis.^16^ A third, which our findings support, is to exploit the dependencies that cells acquire while dormant. Because susceptibility to metabolically induced quiescence is itself subgroup-specific, the choice between these strategies may be dictated by the intrinsic state of the tumour, favouring dependency-directed targeting in contexts where forced activation would be neither selective nor safe.

How DENSpm treatment leads to dsRNA accumulation remains unclear. One possibility is that polyamine depletion alters chromatin accessibility at ERV loci; ATAC-seq across DENSpm-treated GSCs would establish whether this is the case and provide a basis for dissecting the underlying epigenetic mechanism. Such regulatory dynamism would be consistent with the compartment-specific shifts in distal enhancer accessibility seen as brain tumour stem cells progress down a lineage hierarchy.^16^ Alternatively, or in addition, polyamines may act post-transcriptionally to modulate the stability and degradation of dsRNA. A central open question is whether polyamine alteration is a cause or a consequence of the mesenchymal shift toward a more proliferative, quiescence-refractory state. Resolving the relationship between the inflammatory state of GSCs, polyamine remodelling and therapeutic resistance will be essential to understand the dual role of inflammation in both enforcing quiescence and sustaining the mesenchymal state. Long-term culture of DENSpm-sensitive GSCs under putrescine-restricted conditions offers a tractable means to ask whether a pro-inflammatory, DENSpm-resistant state can be selected for over time, while spatial transcriptomics relating polyamine content to transcriptional identity across distinct tumour regions would help connect these states in situ. Although a pro-inflammatory shift may

itself be a necessary step down the stem cell hierarchy toward aggressive more mesenchymal or injury response GSCs, establishing a direct mechanistic link to GSC quiescence would open new therapeutic avenues. Finally, our findings are confined to in vitro systems, and whether disrupting this circuit prevents regrowth in vivo remains to be tested.

Focusing here on metabolic control of stem cell states, we have defined a polyamine-driven, innate immune-mediated program that induces quiescence in GSCs. Engaging this program imposes a growth arrest that spares the stem cell pool, while blocking the lysosomal dependency it creates prevents cells from resuming growth once the arrest is established. Clinically, additional therapies for patients with GBM are urgently needed. Although quiescence itself remains a challenging cancer stem cell property to target, exploiting the dependencies that cells acquire while dormant can constrain the tumor seeding and repopulating fraction by acting on a specific state within the hierarchy. We propose that pairing polyamine-directed treatment with disruption of lysosomal function may find a role in combination therapy for GBM, in the subgroup of tumours whose intrinsic inflammatory tone renders them susceptible. Unlocking metabolically regulated cell state transitions suggests new treatment approaches for limiting the persistence of stem cells at the apex of a neoplastic hierarchy.

## Methods

### Tissue Culture and Reagents

Patient-derived glioblastoma stem cells (GSCs) were cultured as previously described.^50^ Adherent GSC lines (annotated Gxxx) were grown on laminin-coated tissue culture plates in serum-free NeuroCult NS-A Basal Medium (StemCell Technologies) supplemented with NeuroCult Proliferation Supplement (StemCell Technologies), 20 ng/mL EGF (Peprotech), 20 ng/mL bFGF (Peprotech), and 2 μg/mL heparin (Sigma-Aldrich). Cells were passaged using enzymatic dissociation with Accutase (StemCell Technologies). All cultures were maintained at 37°C with 5% CO . Primary patient-derived GBM samples were obtained with informed consent under protocols approved by the Research Ethics Board at The Hospital for Sick Children (SickKids). Human fetal neural stem (hfNSC) cell lines were used as normal CNS stem cell controls grown under identical conditions. All cell lines were routinely screened for mycoplasma contamination.

### Pharmacologic treatments

Pharmacological compounds were reconstituted and stored according to the manufacturer’s instructions; working concentrations are specified in the relevant figure legends. DENSpm (N^1^,N^11^- Diethylnorspermine; (Tocris Bioscience, cat # 0468) was reconstituted in dH_2_O and applied at 10 µM unless otherwise stated, with media. For washout experiments, cells were treated for 7 days, washed in PBS, and returned to drug-free medium. Cytosine arabinoside (Ara-C; Sigma-Aldrich) was used at 1 µM as a comparator cytotoxic agent. Recombinant human IFN-β (Gibco) was applied at 100–1000 units/mL as indicated. Lysosomal/autophagy inhibitors bafilomycin A1 (Sigma-Aldrich, cat # B1793) was applied alone or in combination with DENSpm as indicated at 100nM concentration. Methylated polyamine analogues were obtained from Dr. Tuomo A. Keinänen.^38^ Vehicle-treated cells served as controls throughout.

### NucRed Live/Dead Staining and Senescence Assay

Cell viability was assessed using the NUCLEAR-ID® Red DNA stain (Enzo Life Sciences, cat #ENZ- 52406) according to the manufacturer’s protocol. NucRed-positive (dead) and NucRed-negative (live) populations were distinguished by fluorescence using the IncuCyte ZOOM™ Live Cell Analysis System. Cell viability was calculated from cell images taken every 8 hours using the IncuCyte ZOOM software.

Senescence-associated β-galactosidase activity was assessed after 7 days of DENSpm (10 µM) or control treatment using the CellEvent™ Senescence Green Detection Kit (ThermoFisher Scientific, cat # C10850) according to the manufacturer’s instructions.

### Label-Retaining Experiment

To identify slow-cycling/label-retaining cells, GSCs were transfected with a H2B–GFP reporter (Addgene, cat #11680) and sorted for stable expression. Label-retaining (GFP-high), slow-cycling (GFP- intermediate) and fast-cycling (GFP-low) populations were resolved and cell sorting was performed on either a Beckman Coulter MoFlo or Beckman Coulter MoFlo-XDP cell sorter. Data were analyzed using FlowJo software.

### Flow Cytometry, Fluorescence-Activated Cell Sorting (FACS), and Cell Cycle Analysis

For flow cytometry analysis and FACS, adherent GSCs were dissociated using Accutase (StemCell Technologies), then washed in PBS. Viability was assessed by inclusion of DAPI immediately prior to acquisition. Cell sorting was performed on either a Beckman Coulter MoFlo or Beckman Coulter MoFlo-XDP cell sorter. Doublets were excluded using FSC-A/FSC-H gating; dead cells were excluded with a viability dye. Data were analyzed using FlowJo v10 (BD Biosciences).

For cell cycle analysis, dissociated cells were fixed in ice-cold 70% ethanol overnight at −20°C, washed twice in PBS, and resuspended in propidium iodide (PI) staining solution (50 μg/mL PI, 100 μg/mL RNase A in PBS) for 30 minutes at 37°C protected from light. DNA content was measured by flow cytometry and G0/G1, S, and G2/M fractions were quantified using FlowJo software.

### Proliferation and self-renewal assays

Cell growth was monitored using the IncuCyte ZOOM™ Live Cell Analysis System. Confluency was calculated from cell images taken every 8 hours using the IncuCyte ZOOM software.

For limiting dilution assays (LDA), the GSCs were plated in serial dilutions on non-adherent 96-well plates with six technical replicates per dilution in the media described above for adherent. LDAs were performed by plating cells in suspension in 1:2 serial dilution as follows: 2000, 1000, 500, 250, 125, 63, 31, 16, 8, 4 cells per well. Each cell dilution was plated as six technical replicates, all on the same plate. Each well was replenished with 50 μL of fresh media after 1 week. 7 and 14 days after plating, wells from each plate were scored blindly for the presence of spheres.

For the secondary limiting dilution assay, spheres collected from the primary assay after 14 days. They were mechanically and enzymatically (0.05% Accutase) dissociated followed by filtering and visualisation with Trypan Blue to ensure a single cell suspension. They were then counted and replated for the subsequent limiting dilution assay experiment. The readout at the assay endpoint was the presence or absence of spheres in each well. The fraction of technical replicates positive for the presence of spheres at each dilution was entered into the extreme limiting dilution analysis (ELDA) software (http://bioinf.wehi.edu.au/software/elda/) to determine the estimated sphere-forming frequency along with upper and lower limits (denoted by the error bars) with a 95% confidence interval. Data were tested for inequality in frequency between multiple groups and for adequacy of the single-hit model using the ELDA software.^51^

### Immunofluorescence

Cells were fixed in 4% PFA, permeabilized in 0.5% Triton X-100, blocked in BSA, and incubated with primary antibodies against SOX2 (R&D Systems, cat #AF2018), GFAP (Abcam, cat #ab4674), βIII- tubulin (TUJ1, Sigma-Aldrich, cat #MAB1637), double-stranded RNA (J2, Sigma-Aldrich, cat#MABE1134), IRF7 (Invitrogen, cat #PA5-85724), phospho-IRF7 (Ser477, Cell Signaling Technology cat #12390), and TFEB antibody was generously provided by Dr. John Dick,^46^ followed by Alexa Fluor 680 Donkey anti-Rb IgG (Invitrogen cat #A10043) and DAPI counterstain. Nuclear translocation of phosphoIRF7 was quantified as nuclear/cytoplasmic signal ratio. Images were acquired using a Leica DMi8 Inverted LED Fluorescence Motorized Microscope and analysed using the Leica Application Suite software.

### Protein synthesis (OPP) assay

Global translation was measured by O-propargyl-puromycin (OPP) incorporation. Cells were pulsed with OPP for 60 min, fixed, and the alkyne label detected by the Click-iT™ Plus OPP Alexa Fluor™ 488 Protein Synthesis Assay Kit (Thermo Fisher Scientific, cat #C10456) as per manufacturer’s protocol.

Mean OPP fluorescence was quantified by imaging and quantified using ImageJ software.

### Protein extraction and Western blot

Lysis buffer containing 10 mM Tris-HCl, 0.5% Triton X-100, 150 mM NaCl, 1 mM EDTA, 10 mM MgCl2, 1% sodium dodecyl sulphate, protease inhibitors, and benzonase was used to harvest total protein for western blot. In all, 30 µg of protein was loaded per well. Cell lysates were run on 10% Bis-Tris SDS-PAGE protein gels and transferred onto polyvinylidene difluoride membranes. Primary antibodies used were SAT1 (Cell Signaling Technology, cat #61586), hypusinated eIF5A (Sigma-Aldrich, cat #ABS1064) and total eIF5A (Abcam, cat #ab32443), MAVS (Santa Cruz, cat #sc-166583), histone methylation marks: H3K4me2 (Abcam #ab177793), H3K9me2 (Abcam, cat #ab1220), H3K9me3 (Abcam cat #ab8898), H3K27me3 , H3K36me2 (Epigentek, cat #A-4041). Secondary antibodies used were anti-mouse-HRP 1:10,000 (Millipore Cat# 2774) and IRDye® 800CW Donkey anti-Mouse 1:15,000 (Licor Cat# P/N 925-32212). Membranes were visualized on an Odyssey® CLx Imaging System (LI-COR).

### Lentiviral SAT1 overexpression and MAVS knockdown

SAT1 was overexpressed by cloning into a dox-inducible lentiviral vector Lenti-X™ Tet-One™ Inducible Expression System (Takara, cat #631844). MAVS was depleted using MAVS shRNA (h) Lentiviral Particles (Santa Cruz Biotechnology; sc-75755-V). Lentivirus was produced in HEK293T cells and GSCs transduced (MOI: 10), then selected with puromycin. Knockdown and overexpression efficiency were confirmed by RT-qPCR and Western blot. For rescue experiments, the effect of MAVS knockdown on DENSpm-induced arrest was assessed over a 7-day treatment course.

### PCR

Total RNA was extracted from GSC cultures or FACS-sorted populations was extracted using the RNeasy Plus Mini Kit (Qiagen cat# 74134), reverse-transcribed using the Sensiscript RT Kit (Qiagen cat# 205213), and quantified by RT-qPCR(Qiagen) with on-column DNase I digestion. For quantitative RT-PCR, reactions were run in technical triplicate using SYBR Green Master Mix (Applied Biosystems) on a QuantStudio 6 or equivalent instrument. Relative expression was calculated by the 2^(−ΔΔCt) method normalized to ACTNB and RPLPO. Primers used can be found in Supplementary Table 10.

### Metabolomics

GSCs and hfNSCs, untreated or treated with DENSpm, were washed twice in ice-cold PBS and snap-frozen as dry pellets (8 × 10^6^ cells; n = 3 per condition). Untargeted metabolomic profiling was performed by Metabolon Inc. (Morrisville, NC, USA), as described previously.^52^ Briefly, metabolites were extracted with methanol on an automated liquid handler, and the extracts were split for analysis by four UPLC– MS/MS methods: two reverse-phase positive-ion, one reverse-phase negative-ion and one HILIC negative-ion. Analysis used a Waters ACQUITY UPLC coupled to Thermo Q Exactive high-resolution Orbitrap mass spectrometers (1, 2). Metabolites were identified by matching retention index, accurate mass (±10 ppm) and MS/MS spectra to an in-house library of authentic standards (MSI level 1).^52^

Relative levels of putrescine, spermidine, spermine and SAM were calculated from peak areas, normalized, and expressed relative to untreated controls. Representative data are shown for G549, G564 and G583.

### DNA methylation profiling

Genomic DNA from DENSpm-treated and control GSCs was bisulfite-converted and profiled on Infinium MethylationEPIC BeadChip arrays (Illumina) as previously described (Guilhamon et al., 2021).^39^ Briefly, to process and analyze the data the R package ChAMP v2.6.4^53^ was used. For the copy number analysis, the raw IDAT files were uploaded to the MNP tool,^54^ which compares the copy number profile estimated from the probe intensities on the methylation array to the distribution observed across thousands of brain tumors in its database.

### Bulk RNA sequencing and derived signatures

#### Bulk RNA Sequencing

Total RNA was extracted from FACS-sorted or bulk GSC cultures using the RNeasy Plus Mini Kit (Qiagen Cat# 74134) with on-column DNase digestion using the gDNA Eliminator Columns. RNAseq libraries were constructed using the NEB Ultra II Directional polyA mRNA Library Prep Kit (NEB Cat# E7765) with on-column DNase I digestion. RNA quality was confirmed by Agilent Bioanalyzer (RIN ≥ 7 required). Libraries were prepared from the TruSeq Stranded mRNA Library Prep Kit (Illumina), per the manufacturer’s protocols. Libraries were sequenced (100 bp, paired-end) on an Illumina HiSeq or NovaSeq at The Centre for Applied Genomics (TCAG), SickKids. FASTQ quality control was performed with FastQC and trimming with Trimmomatic. Reads were aligned to hg38 using STAR and gene-level counts derived with featureCounts. Differential expression was assessed with DESeq2 (adjusted p < 0.05; log FC > 1). Gene set enrichment analysis (GSEA) was performed with the fgsea R package against hallmark and curated gene sets from MSigDB.

#### The following signatures were derived and/or applied

DENSpm_Q signature: genes significantly upregulated after one week of DENSpm across responsive lines; applied to scRNAseq and bulk data as a per-cell/per-sample quiescence score.

DENSpm response-subgroup signatures (Non-reversible, Reversible, Resistant): genes exclusively upregulated by DENSpm within each response group; applied across bulk, single-cell and spatial datasets. Published quiescence signatures: Neural.G0 (neural stem cell quiescence)^34^ and the GBM quiescence signature Q118,^6^ used to benchmark induced and basal quiescence.

TFEB-induced quiescence signature (TFEB_HSC): correlated with SAT1 expression in glioma cohorts. Single-cell RNA-seq analysis (reanalysis of published data).

Previously published patient-derived GSC scRNA-seq cohort of Richards et al. (*Nature Cancer* 2021; 10x Genomics; GEO accession: GSE117529)^24^ was re-analyzed for this study. Cells were scored for cell-cycle phase using Seurat CellCycleScoring with canonical G1/S and G2/M gene sets. Quiescence (Neural.G0, Q118), DENSpm_Q, and response-subgroup signatures were scored per cell. Genes enriched in the non-cycling compartment were identified by differential expression and summarized by Gene Ontology enrichment analysis.

### ATAC-seq / transposable-element accessibility (reanalysis of published data)

Chromatin accessibility was assessed by reanalysis of the previously published GSC ATAC-seq cohort of Guilhamon et al. (*eLife* 2021; GEO accession: GSE109399)^39^; no new ATAC-seq was generated here.

Processed peaks were used to quantify accessibility across transposable-element families; mean accessibility per TE subfamily was compared across endogenous retroviral (ERV) elements against other TE classes across untreated GSCs.

## Public cohort and survival analyses

SAT1 expression was related to overall survival in GBM and low-grade glioma using TCGA, with patients stratified by *SAT1* level and compared by Kaplan-Meier analysis. *SAT1* expression was correlated with the TFEB_HSC signature in TCGA GBM.

## Quantification and statistical analysis

Three independent biological replicates were performed for each experiment unless otherwise stated. Data are presented as mean ± SD. Comparisons between two groups used two-tailed Student’s t-test Mann– Whitney; correlations used Spearman/Pearson as defined in the figure legend. Statistical significance was defined as *p* < 0.05, and analyses were performed in GraphPad Prism.

## Z-Scoring

For comparative visualization of gene expression or metabolomic data across samples and conditions, values were z-score normalized. For each gene or feature, the mean was subtracted and divided by the standard deviation across the group being compared. Z-scoring was applied to log-transformed and/or variance-stabilized matrices, as appropriate to the data type (bulk RNA-seq, scRNA-seq, microarray, or metabolomics). Normalization was performed using the scale() function in R, and results were visualized as heatmaps using pheatmap or ComplexHeatmap packages.

## Data and code availability

Newly generated bulk RNA-seq, and DNA methylation array data have been deposited at [GEO xxxxxxxxx].

## Supplementary Figure Legends

**Supplementary Figure 1.** : **(A)** Cell cycle scatter plot highlighting selected non-cycling cells in green. **(B)** Scatter plots of quiescence signatures (Q118 and Neural.G0) and ridge plots showing the distribution of **(C)** Q118 and **(D)** Neural.G0 scores according to cell cycle states. **(E)** Representative flow cytometry figures illustrating the presence of a quiescent G0 population in GSCs (G564). **(F)** Graph showing doubling time of the three sorted populations (P1 to P3) after initial plating post-sorting (P0) (G549).

**Supplementary Figure 2.** :**(A)** G0vsG1 gene ontology treemap and **(B)** Violin plot of SAT1 expression according to cycling state across GSCs. **(C)** DENSpm dose curves based on Alamar blue fluorescence 14 days post-treatment. **(D)** Individual and pooled replicated LDA experiments on GSCs while treated with DENSpm and **(E)** LDA plated post DENSpm treatment. **(F)** Volcano plot of differential expression analysis on DENSpm treated GSCs. **(G)** Ridge plot of DENSpm quiescence score (DENSpm_Q) according to cell cycle phases in scRNAseq data and **(H)** cell cycle scatter plot colored based on DENSpm_Q. **(I)** SAT1 ridge plot of cells segregated according to DENSpm_Q.

**Supplementary Figure 3.** :**(A)** Representative western blot showing hypusination of EIF5A upon DENSpm treatment and SAT1 overexpression (G549). **(B)** scatter plot of untreated GSCs according to polyamine content, SAM level, and global DNA methylation. **(C)** Bar graph depicting SAM level in GSCs following DENSpm treatment in G549, G564, and G583. **(D)** Bar graph showing the number of methylated probes and **(E)** histogram of deltaBeta values based on Infinium Methylation EPIC array performed on a representative cohort of GSCs treated with DENSpm. **(F)** Pearson’s correlation heatmap of median absolute deviation of the top 1 percent most variable probes. **(G)** Representative Western blot of various histone methylation marks in four GSCs (G567, G564, G549 and G523) treated with vehicle control or DENSpm. **(H)** Graph representing activated and suppressed pathways identified through GSEA in GSCs upon SAT1 overexpression and **(I)** Emap plot. **(J)** KEGG pathview showing regulation of genes in GSCs treated with DENSpm in the cell cycle and **(K)** dsRNA detection pathway maps.

**Supplementary Figure 4.** :**(A)** Representative images of dsRNA immunolabelling (J2 antibody) in GSCs treated for a week with DENSpm and after DENSpm wash. **(B)** Bar graph of mean accessibility of ERVs. **(C)** Box graph of mean accessibility of mature and immature TE in GSCs.^35^ **(D)** Scatter plot of GSCs basal ERVs z-score and IFNβ. **(E)** Scatter plot of GSCs basal ERVs z-score and ISG. **(F)** bar graph of SAT1 expression in GSCs measured by RT-qPCR. **(G)** Representative western blot of SAT1 in a representative GSC cohort.

**Supplementary Figure 5.** :**(A)** Growth and cytotoxicity curves in GSCs treated with DENSpm or Ara-C. **(B–C)** Bar graph of IFNs (B), and ISGs (C) regulation after a 7-day DENSpm treatment and measured by RT-qPCR in GSCs and hfNSCs segregated according to response subgroups. **(D)** Table of GSCs classification. **(E)** GSEA enrichment graphs and **(F)** dsRNA detection pathview maps comparing resistant GSCs with non-reversible or reversible GSCs (untreated). **(G)** GSEA enrichment graphs and **(H)** dsRNA detection pathview maps looking at the impact of DENSpm in the three DENSpm response groups.

**Supplementary Figure 6.** :**(A)** Venn diagram representing uniquely upregulated genes in non-reversible, reversible, and resistant GSCs. **(B)** Scatter plots of a large cohort of GSCs scattered based on quiescence signatures (Q118 and Neural.G0) and colored according to Non-reversible, Reversible and Resistant signature scores. **(C)** Violin plots of unique response scores in GSCs grouped based on quiescence scores. **(D)** Scatter plots of a large cohort of GSCs developmental and injury response scores and colored according to Non-reversible, Reversible and Resistant signature scores. **(E)** Violin plots of unique response scores in GSCs grouped as developmental or injury response GSCs. **(F)** Violin and ridge plots of cycling, quiescence, and developmental to injury response scores in distinct GSCs populations identified based on our response subgroup specific gene signatures. **(G)** Box graphs of expression of putative polyamine lysosomal transporter and SAT1 upon DENSpm treatment (log2 counts from RNAseq data).

## Notes

### Competing Interest Statement

The authors have declared no competing interest.

