## Supplementary Figures for "Metabolic rewiring induces a viral mimicry-mediated quiescence in glioblastoma"

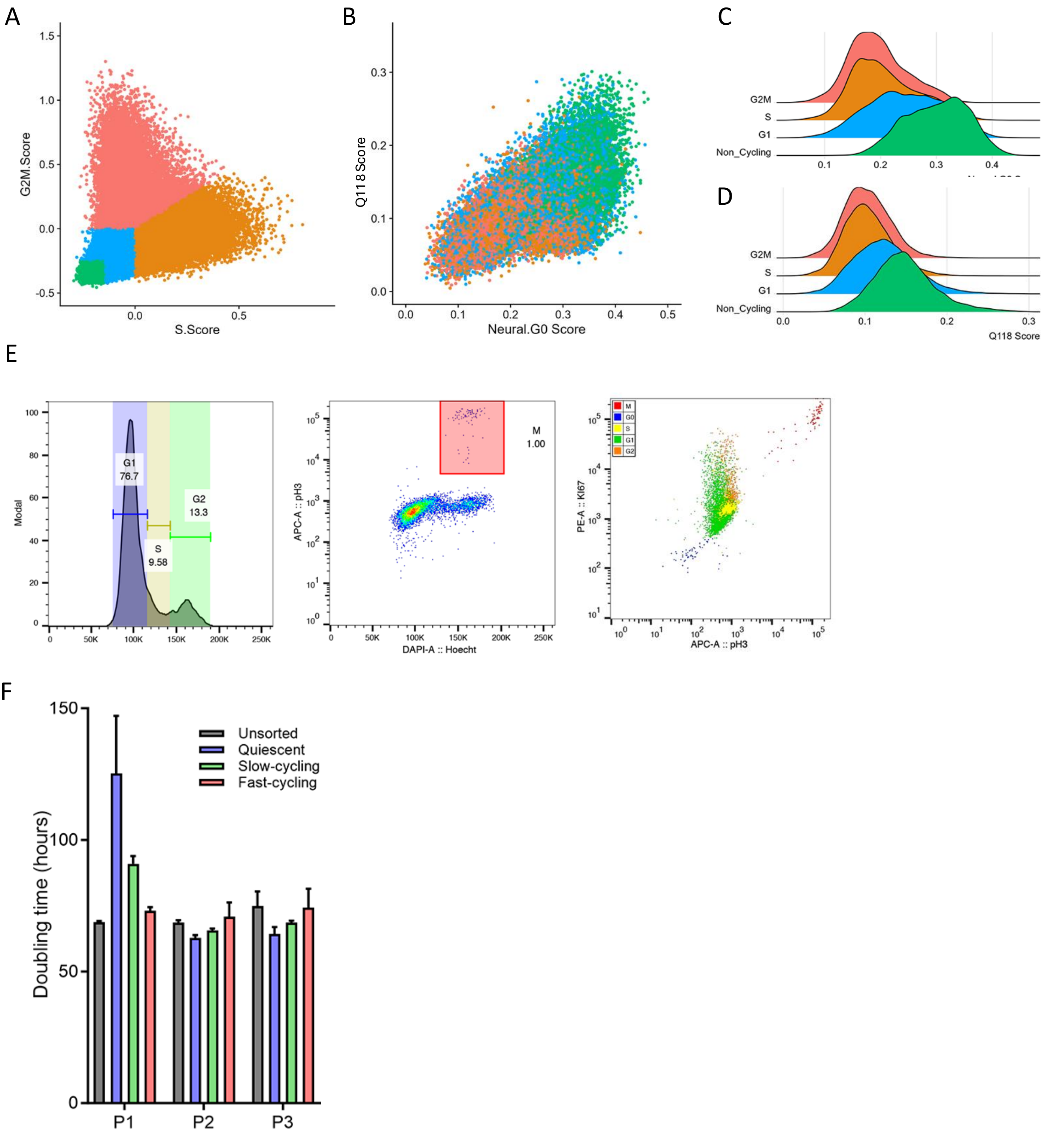


Supplementary Fig. 1

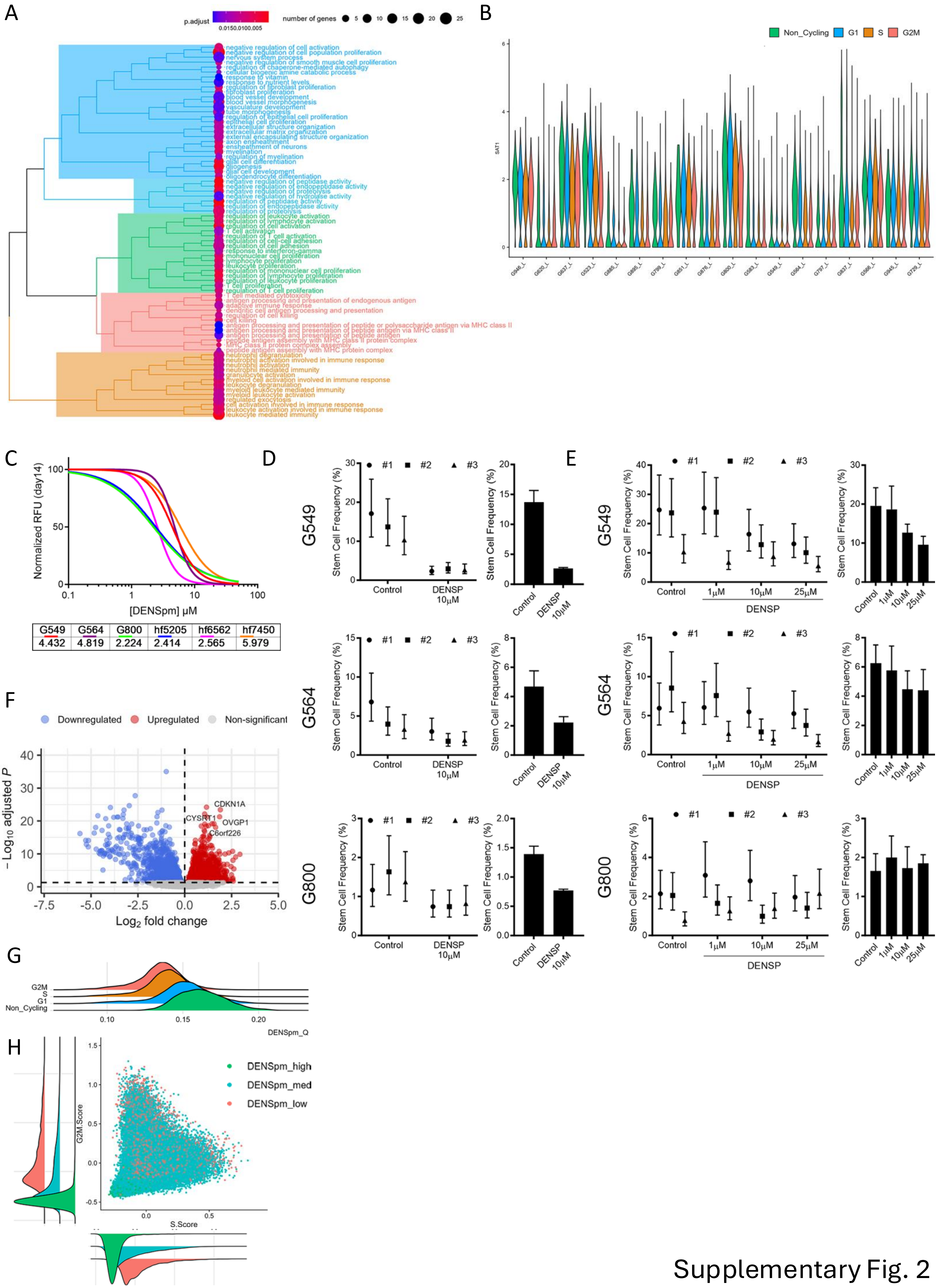


Supplementary Fig. 2

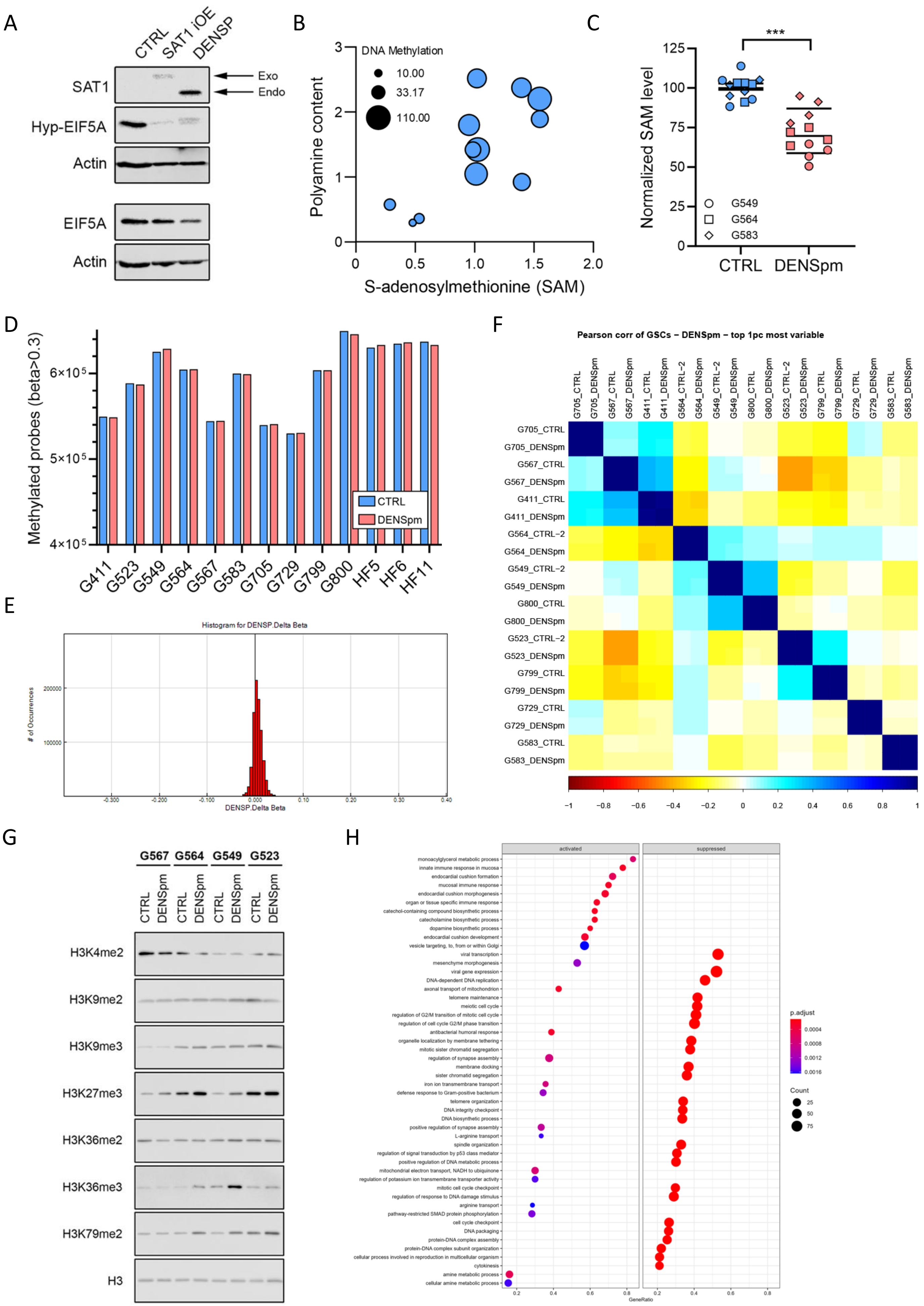

A
CTRL
SAT1 iOE
DENSP
SAT1
Hyp-EIF5A
Actin
EIF5A
Actin
Exo
Endo
B
DNA Methylation
10.00
33.17
110.00
Polyamine content
S-adenosylmethionine (SAM)
C
***
Normalized SAM level
G549
G564
G583
CTRL
DENSpm
D
Methylated probes (beta>0.3)
CTRL
DENSpm
G411
G523
G549
G564
G567
G583
G705
G729
G799
G800
HF5
HF6
HF11
E
Histogram for DENSP.Delta Beta
### of Occurrences
DENSP.Delta Beta
F
Pearson corr of GSCs – DENSpm – top 1pc most variable
G705_CTRL
G705_DENSpm
G567_CTRL
G567_DENSpm
G411_CTRL
G411_DENSpm
G564_CTRL–2
G564_DENSpm
G549_CTRL–2
G549_DENSpm
G800_CTRL
G800_DENSpm
G523_CTRL–2
G523_DENSpm
G799_CTRL
G799_DENSpm
G729_CTRL
G729_DENSpm
G583_CTRL
G583_DENSpm
G
G567
G564
G549
G523
CTRL
DENSpm
H3K4me2
H3K9me2
H3K9me3
H3K27me3
H3K36me2
H3K36me3
H3K79me2
H3
H
activated
suppressed
monoacylglycerol metabolic process
innate immune response in mucosa
endocardial cushion formation
mucosal immune response
endocardial cushion morphogenesis
organ or tissue specific immune response
catechol-containing compound biosynthetic process
catecholamine biosynthetic process
dopamine biosynthetic process
endocardial cushion development
vesicle targeting, to, from or within Golgi
viral transcription
mesenchyme morphogenesis
viral gene expression
DNA-dependent DNA replication
axonal transport of mitochondrion
telomere maintenance
meiotic cell cycle
regulation of G2/M transition of mitotic cell cycle
regulation of cell cycle G2/M phase transition
antibacterial humoral response
organelle localization by membrane tethering
mitotic sister chromatid segregation
regulation of synapse assembly
membrane docking
sister chromatid segregation
iron ion transmembrane transport
defense response to Gram-positive bacterium
telomere organization
DNA integrity checkpoint
DNA biosynthetic process
positive regulation of synapse assembly
L-arginine transport
spindle organization
regulation of signal transduction by p53 class mediator
positive regulation of DNA metabolic process
mitochondrial electron transport, NADH to ubiquinone
regulation of potassium ion transmembrane transporter activity
mitotic cell cycle checkpoint
regulation of response to DNA damage stimulus
arginine transport
pathway-restricted SMAD protein phosphorylation
cell cycle checkpoint
DNA packaging
protein-DNA complex assembly
protein-DNA complex subunit organization
cellular process involved in reproduction in multicellular organism
cytokinesis
amine metabolic process
cellular amine metabolic process
GeneRatio
p.adjust
Count

I

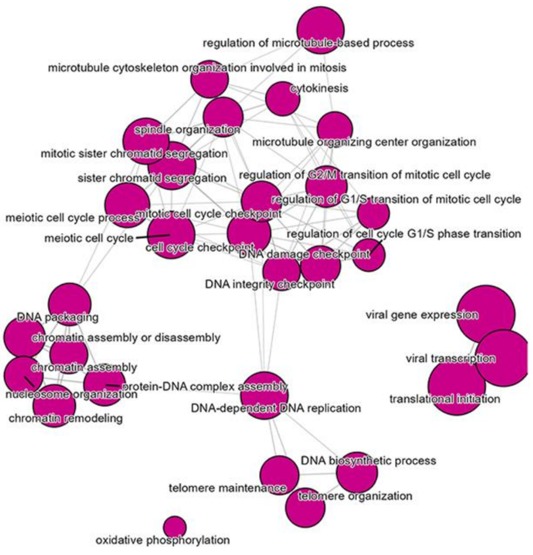


J

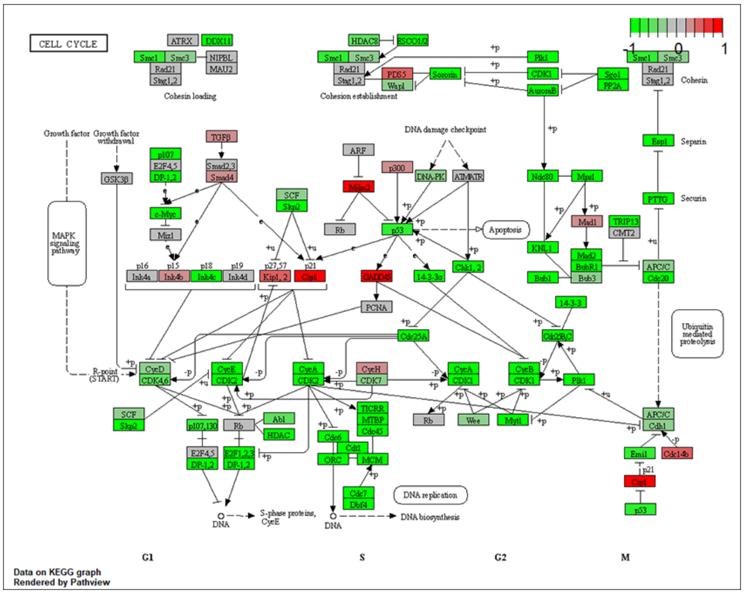


K

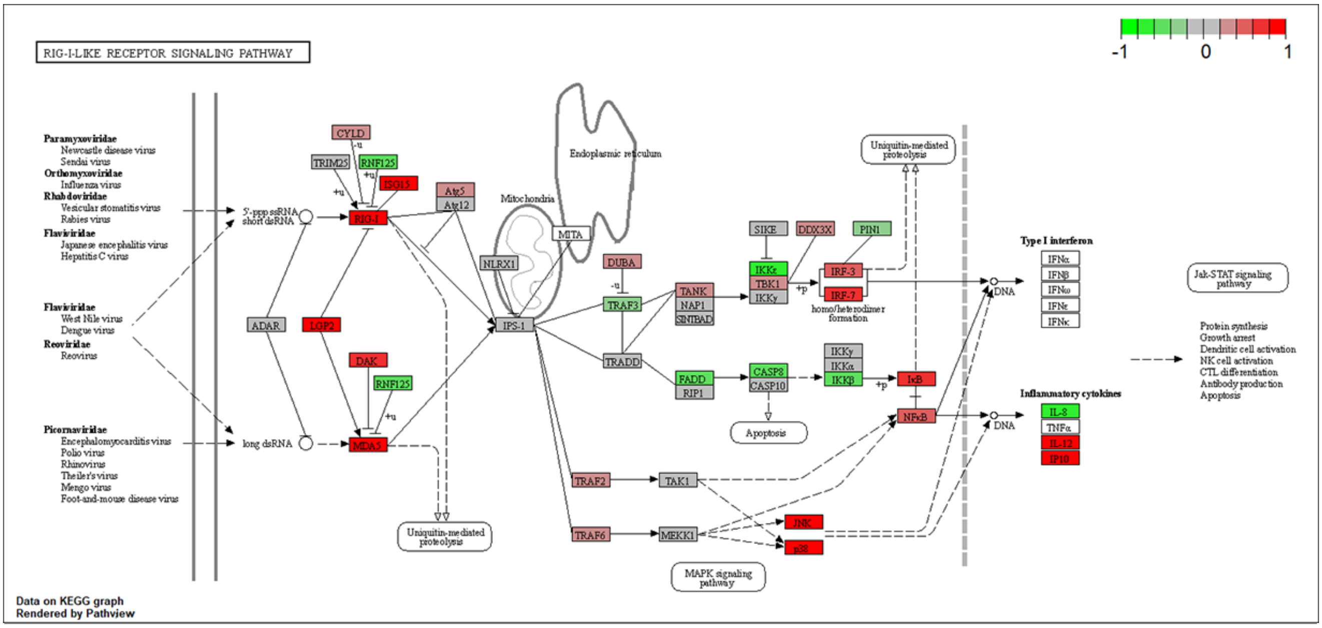


Supplementary Fig. 3

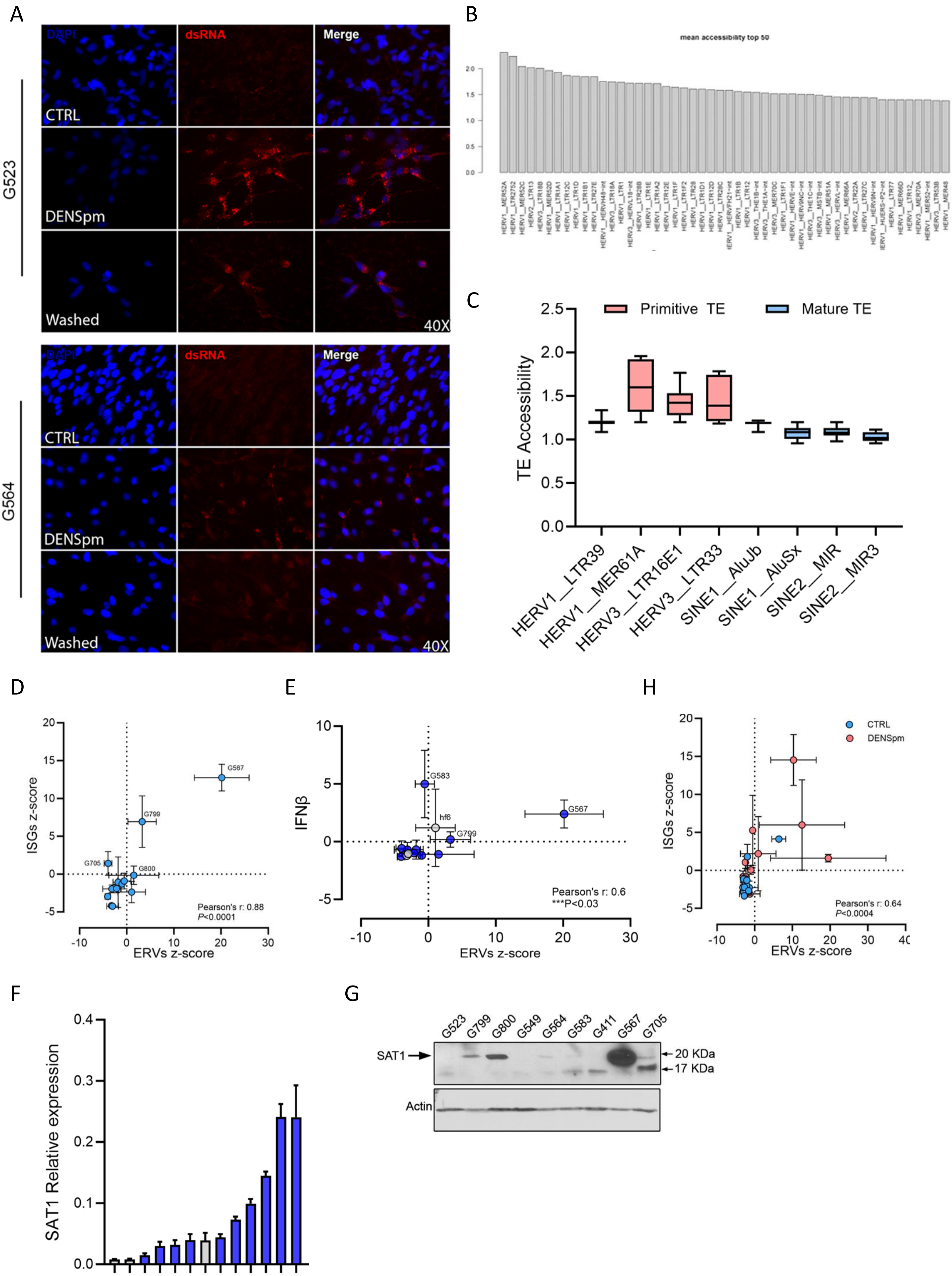


Supplementary Fig. 4

A
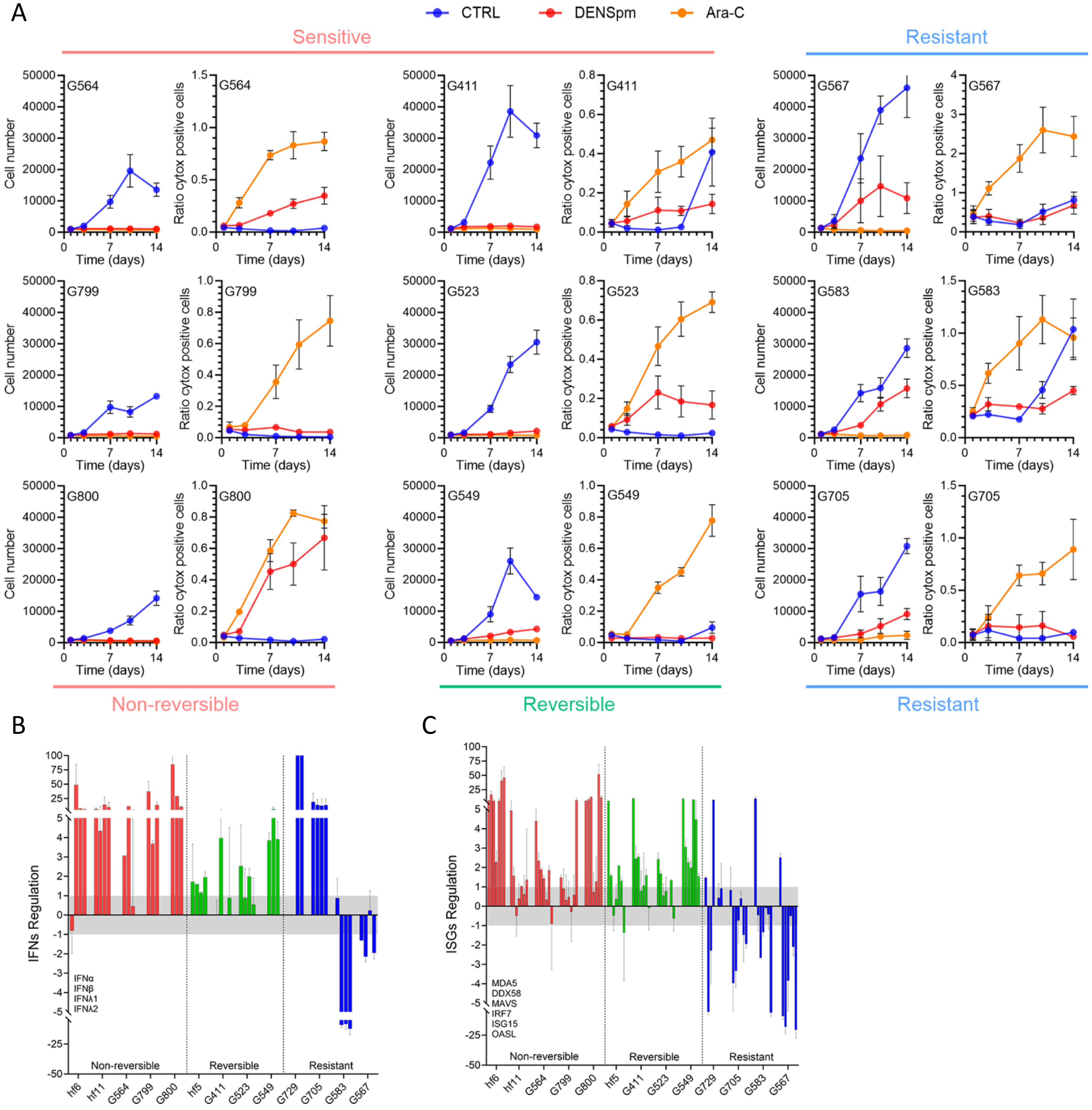


D

| | Subgroups | | | |
|---|---|---|---|---|
| **Lines** | **Cell Type** | **GSC groups** (Guilhamon P, et al.) | **GSC groups** (Richards L, et al.) | **DENSpm response** |
| **Hf5** | **hfNSCs** | n.a | n.a | Reversible |
| **Hf6** | | | | Non-reversible |
| **Hf11** | | | | Non-reversible |
| **G523** | **GSCs** | Constructive | Developmental | Reversible |
| **G799** | | | | Non-reversible |
| **G800** | | | | Non-reversible |
| **G549** | | Reactive | Injury response | Reversible |
| **G564** | | | | Non-reversible |
| **G583** | | | | Resistant |
| **G729** | | | | Resistant |
| **G411** | | Invasive | | Reversible |
| **G567** | | | | Resistant |
| **G705** | | | | Resistant |

E

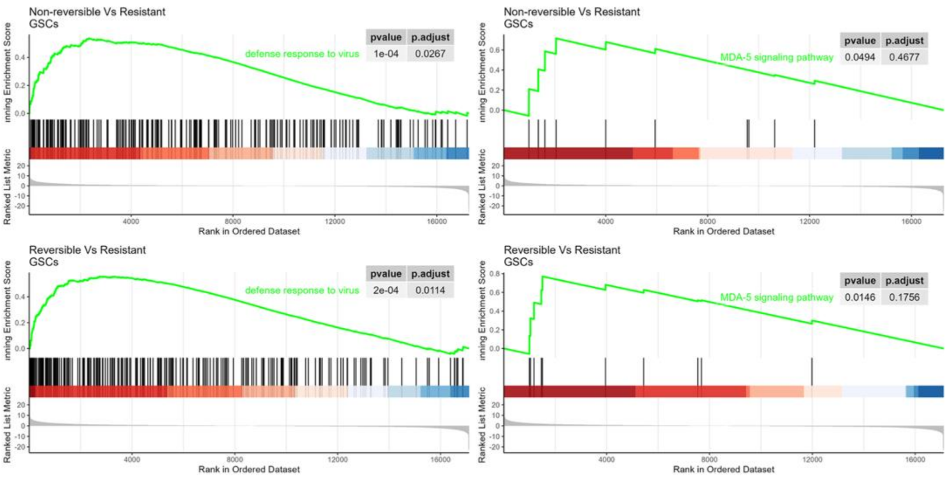


F

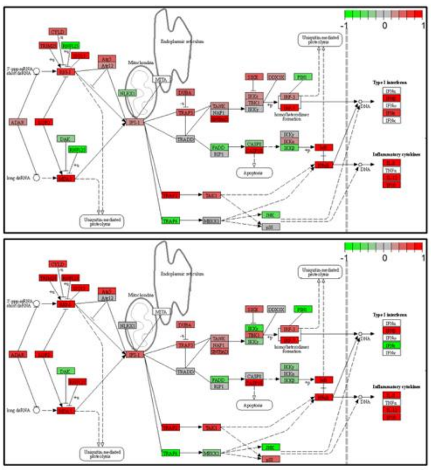

G

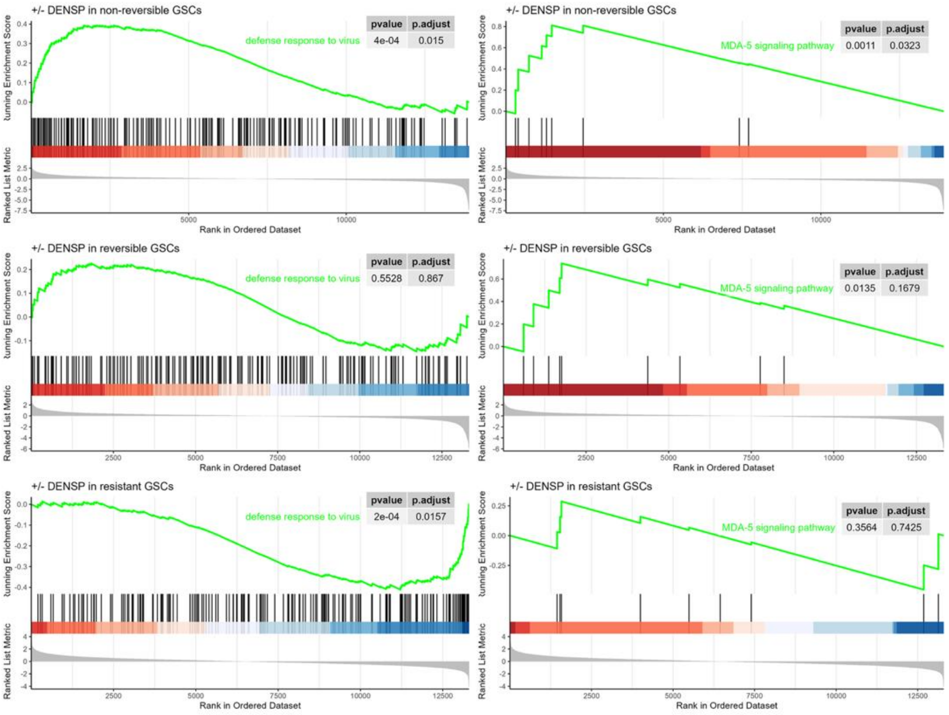


H

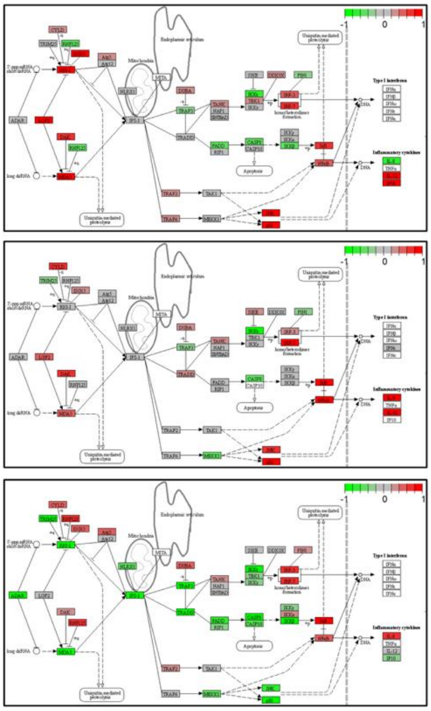

Supplementary Fig. 5

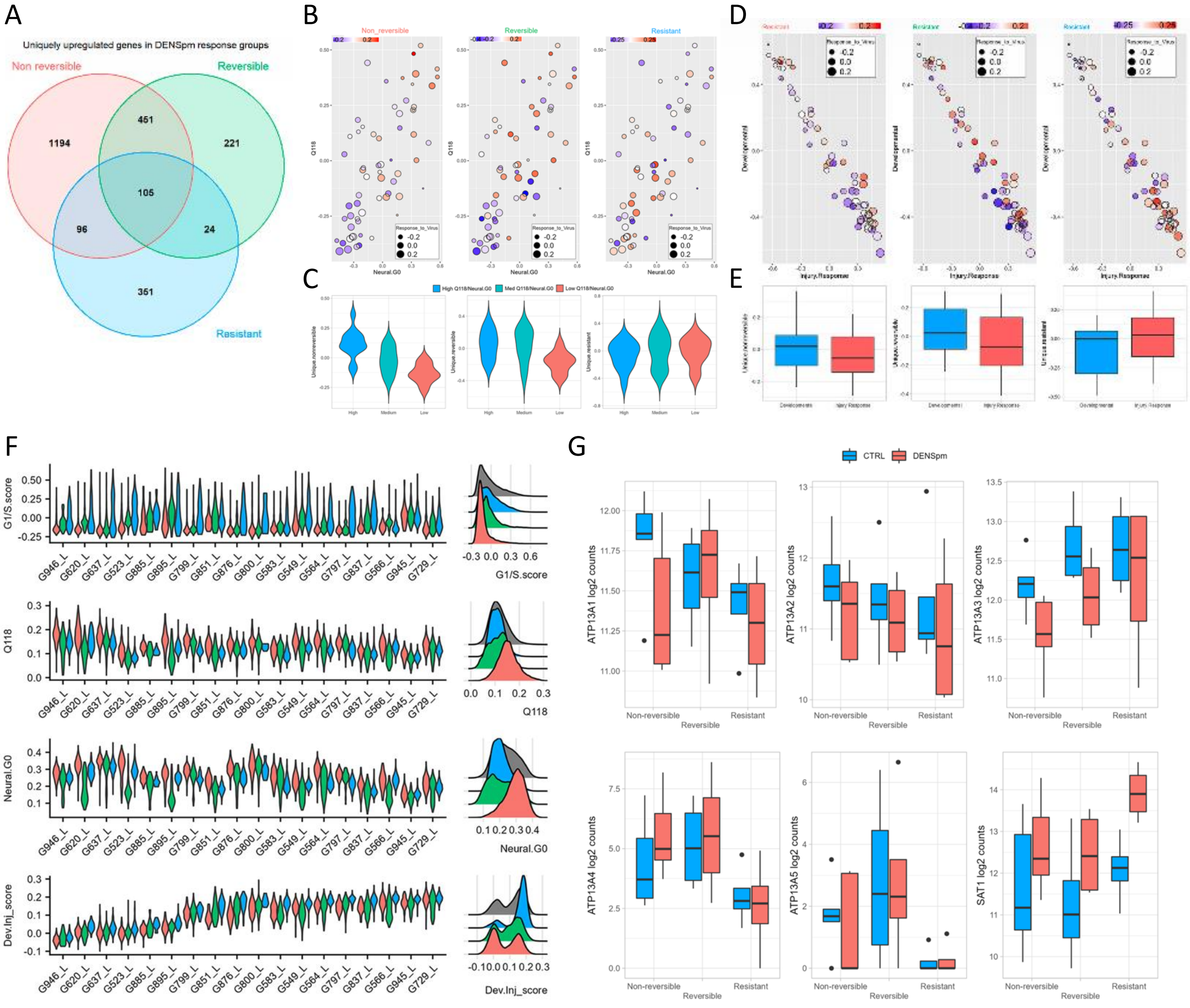


Supplementary Fig. 6
